# Host species identity shapes the diversity and structure of insect microbiota

**DOI:** 10.1101/2021.07.19.452888

**Authors:** Antonino Malacrinò

## Abstract

As for most of the life that inhabits our planet, microorganisms play an essential role in the fitness of insects, including nutrition, reproduction, defence, and many other functions. More recently, we assisted in an exponential growth of studies describing the taxonomical composition of bacterial communities across insects’ phylogeny. However, there is still an outstanding question that needs to be answered: which factors contribute most to shape insects’ microbiomes? This study tries to find an answer to this question by taking advantage of publicly available sequencing data and reanalysing over 4,000 samples of insect-associated bacterial communities under a common framework. Results suggest that insect taxonomy has a wider impact on the structure and diversity of their associated microbial communities than the other factors considered (diet, sex, life stage, sample origin, and treatment). However, when specifically testing for signatures of co-diversification of insect species and their microbiota, analyses found weak support for this, suggesting that while insect species strongly drive the structure and diversity of insect microbiota, the diversification of those microbial communities did not follow their host’s phylogeny. Furthermore, a parallel survey of the literature highlights several methodological limitations that need to be considered in future research endeavours.

## 1 Introduction

Insects are one of the most successful and diverse groups of organisms. Most of the natural and agricultural systems depend on or are heavily influenced by insects (Losey and Vaughan, 2006; Yang and Gratton, 2014). At the same time, microorganisms play an essential role in insect biology, development, fitness, and lifestyle (Douglas, 2009; Colman et al., 2012; Jones et al., 2013; Yun et al., 2014; Douglas, 2015; Malacrinò, 2018). We have known for decades that microorganisms are essential associates for their host, and more recently we became aware that microbial communities – as a whole – can influence insects’ lifestyle (Douglas, 2015). However, little is still known about the factors that influence the structure and diversity of insect-associated microbial communities, and their relative magnitude.

The influence of microorganisms on their host insects is striking, spanning from nutritional support to modifications of insects’ reproductive behavior. Several recent reviews do an excellent job in describing the span of such interactions and the readers can refer to them for deeper insights (Dillon and Dillon, 2004; Lewis and Lizé, 2015; Douglas, 2015; Gurung et al., 2019). Previous studies attempted to understand the drivers of insect-associated microbial communities. A first meta-analysis by Colman et al. (2012) compared the gut microbial communities of 62 insect species using data from clone librarybased studies. Their major finding is that insect microbiota is mostly shaped by insect taxonomy and secondarily by insect diet, however, this study was limited in the sequencing depth for each insect taxon. Using high-throughput sequencing, Jones et al. (2013) tested the effect of insect taxonomy and diet on 39 insect species (whole insects) collected focusing on a restricted timespan and geographic area to decrease the variability due to geography and seasonality. Similarly to Colman et al. (2012), they found that both insect taxonomy and diet have a significant influence on insect-associated microbial communities, but the effect driven by insect taxonomy was stronger. Yun et al. (2014) expended by analysing a wider set of 218 species across 21 insect orders, and found that gut microbial diversity varies across insect taxa, diet, and life stage, but they did not test the contribution of each factor to the overall microbiota diversity and structure. While there is a wide agreement between these studies, there is still a need to synthesise the effect of a wider set of factors on the diversity and structure of insect-associated microbial communities. Furthermore, none of these studies on a large dataset tested whether a co-diversification (phylosymbiosis) between the host and their associated microbiota exists.

An advantage of the era of big data is the enormous amount of publicly available information. This study takes advantage of public data, reanalysing over 4,000 samples of insect-associated bacterial communities under a common framework, and expanding on previous research to cover a wider set of insect taxa at a wider depth and to test a larger set of factors that might influence insect-associated microbial communities. First, this study provides a survey of the current status of microbiota research in insect science to highlight methodological issues and research biases that need to be addressed for an in-depth understanding of insect microbial communities. Second, the publicly available data is reanalysed and compared to test which factor (phylogeny, diet, sex, life stage, sample origin, or treatment) has an impact on the structure and diversity of insect microbiota and to quantify the magnitude of these effects. Third, this work tests the effect of each of those factors on the abundance of different bacterial taxa. Then, on the basis of the wide effect driven by insect taxonomy on insect microbial diversity (also supported by Colman et al. 2012; Jones et al. 2013; Yun et al. 2014), this study tests whether there is a relationship between the diversity of the insect-associated bacterial community and the insect phylogeny.

## 2 Methods

### 2.1 Data selection and collection

The search of the term *‘insect AND (microbiota OR microbiome)’* on Web of Science Core Collection (year range 2010-2020) provided 1,169 records (April, 11 2020). This was implemented with a search on Google Scholar using the keywords *‘[insect order] (microbiota OR microbiome)’* to guarantee the completeness of the dataset, yielding further 26 studies. All records were further filtered to keep only research articles and remove all those papers not performing high-throughput sequencing on 16S rRNA amplicons from insect specimens, yielding 246 studies. This group was further filtered in an effort to create a reliable dataset to be analysed, focusing on studies: (i) with publicly available data deposited on SRA databases, providing an easily replicable pipeline; (ii) that used the Illumina sequencing technology, avoiding the issues related to comparing data from different sequencing platforms; (iii) that used the V3-V4 portions of 16S rRNA gene as a marker, ensuring unbiased OTU clustering. Studies not reporting the used primer pairs, with an unclear distinction between samples when multiple markers were used, without at least 3 biological replicates, and with non-demultiplexed data were discarded from the analysis. This process yielded 91 studies and 7,079 samples (Fig. 1). Each study and sample was assigned a specific ID number to allow to trace back information: the BioProject nr. was used as study ID, the SRA sample nr. was used as sample ID. Each study was searched to collect the following metadata: DNA extraction method, amplified region of the 16S rRNA gene, primer pairs used, sequencing technology and platform, database where the data was deposited, insect species (single OR multiple). The following metadata was collected for each sample: insect species ID, sex, life stage, sample origin (field OR laboratory), diet, tissue, sample treatment (control OR treatment, regardless of the nature of the treatment). Data were then downloaded using SRA Toolkit 2.10.4. A list of all studies included in the meta-analysis is reported in Supplementary Material 1, and a summary is reported in Supplementary Material 2 and Supplementary Material 3.

**Figure 1.**
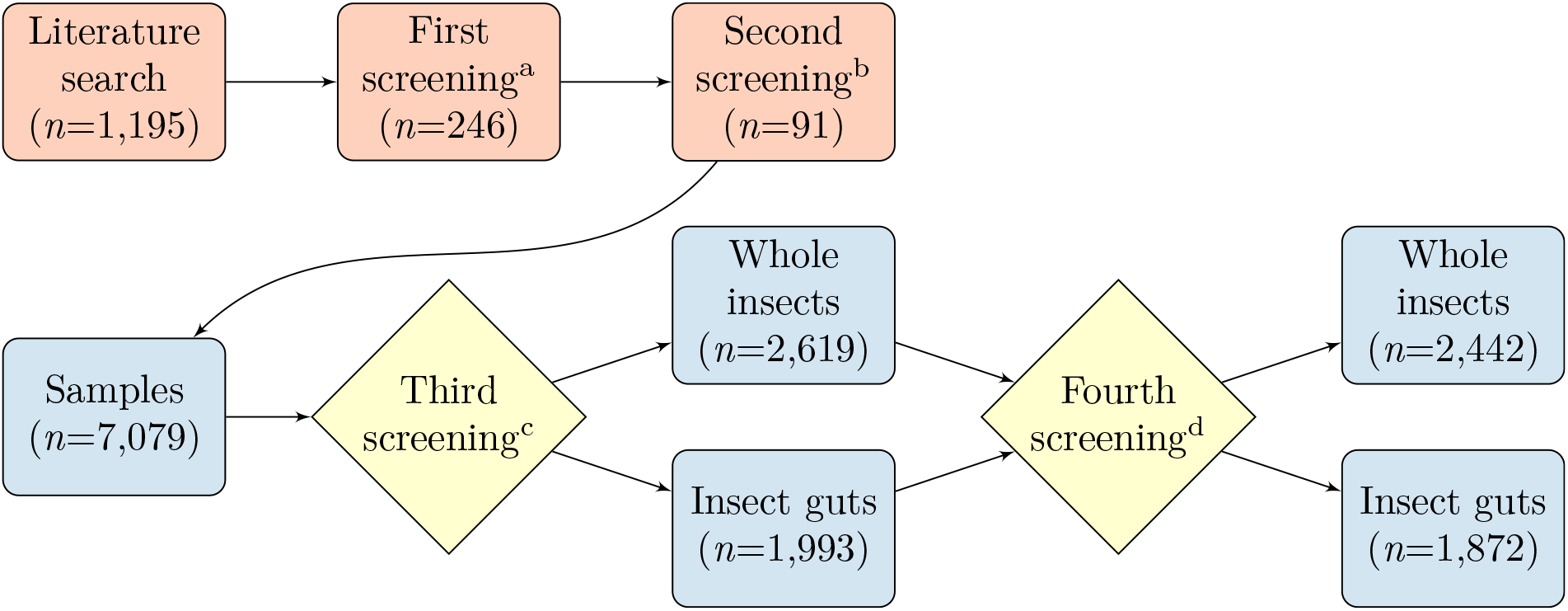
Flow chart showing the selection workflow. The upper row (red) shows the selection of studies, which were screened (a) to consider those relevant for this meta-analysis and (b) to remove those with data not uploaded on SRA, not generated using Illumina technology, not using the V3-V4 portion of the 16S rRNA gene as marker, or with other issues (see Methods). The lower row (blue) shows the selection of samples, which were screened (c) to discard those with <5,000 reads and with insect tissues analysed in less than 3 studies. This lead to consider only two categories: whole insects (both surface sterilized or not) and insect guts. These two groups were further filtered (d) to discard samples with less than 3 species within each order. In the upper row *n* represent the number of studies, while in the lower row *n* represent the number of samples.

### 2.2 Data processing and analysis

Paired-end reads were merged using FLASH 1.2.11 (Magoč and Salzberg, 2011) and data were processed using VSEARCH 2.14.2 (Rognes et al., 2016) with default parameters (including quality filtering, OTU binning, and chimera detection). Taxonomy for representative sequences was determined by querying against the SILVA database v132 (Quast et al., 2012). Data analysis was performed using R statistical software 3.5 (R Core Team, 2020) with the packages *phyloseq* (McMurdie and Holmes, 2013) and *vegan* (Dixon, 2003). In order to test whether our results are influenced by computing OTUs instead of ASVs, all the analyses below have been repeated after processing the data with the DADA2 pipeline (Callahan et al., 2016) and using SILVA v138 for taxonomic assignment, showing that the main results still hold regardless the approach used (Supplementary Material 4).

The initial bulk of 7,079 samples was filtered (Fig. 1) to remove those with <5,000 reads and the OTUs identified in less than 50 samples (~1% of total samples). Reads identified as chloroplast were removed. A summary of all samples included in the meta-analysis is reported in Supplementary Material 3. According to Tab. S3.7 (Supplementary Material 3), most of the studies focused on whole insects or their guts, and just a few analysed other organs. Thus, discarding those tissues analysed in less than 3 studies, this work focuses on broad categories: whole insects (surface sterilised or not, 43 studies and 2,619 samples) and insect guts (41 studies, 1,993 samples). These samples were further filtered to exclude those insect orders with less than 3 different species, yielding 2,442 samples for the group of whole insects (36 studies, 135 species, 5 orders: Hemiptera, Coleoptera, Lepidoptera, Diptera, and Hymenoptera) and 1,872 samples for the group of insect guts (36 studies, 90 species, 7 orders: Odonata, Blattodea, Hemiptera, Coleoptera, Lepidoptera, Diptera, and Hymenoptera). As first step, this study tested the influence of different factors (i.e., insect species, sex, life stage, sample origin, diet, and treatment) on the structure and diversity of insect microbiota. Below, ‘‘structure” refers to the multivariate composition of microbial communities considering both their taxonomical composition and the abundance of the different taxa, while “diversity” refers to a univariate metric (i.e., Shannon index) that summarises the within-sample diversity of that specific microbial community.

A multivariate approach was used to explore the effects of insect species, sex, life stage, sample origin, diet, and treatment on the structure of insect microbiota. Distances between pairs of samples were calculated separately for whole insects and insect guts in two ways: using a Bray-Curtis distance matrix to account for the relative abundance of each OTU, and a Jaccard distance matrix to account only for presence/absence. Each matrix was then tested using PERMANOVA (999 permutations stratified at the level of single study, function *adonis2)* to infer the impact of each factor on the structure of insect microbiota and to estimate the amount of variation explained (formula: *insect_species* + *sex* + *life_stage* + *diet* + *treatment* + *origin).*

The effects of insect species, sex, life stage, sample origin, diet, and treatment on the diversity of insect microbiota were studied using the Shannon diversity index. A linear mixed-effects model was fit separately for whole insects and insect guts including all groups above as fixed factors and the study ID as a random factor (formula: *insect_species* + *sex* + *life_stage* + *diet* + *treatment* + *origin* + (1|*study_ID*)). Models were fit using the *lmer()* function under the lme4 package (Bates et al., 2014), the package *emmeans* was used to infer pairwise contrasts (corrected using False Discovery Rate), and the package *MuMIn* was used to estimate the R^2^ values (Bartoń, 2009) individually for each factor.

When testing the influence of different factors on the diversity and structure of insect microbiota, results suggest that insect species is the one explaining most of the variation (see Results below). To further dissect this result, microbial data were further processed together with an ultrametric phylogenetic tree of insect species obtained from TimeTree (Kumar et al., 2017, accessed on May, 25th 2021). Given that phylogenetic information was not available for all insect species, this analysis was run on a subset of 42 species and just focusing on insect guts. The influence of host phylogeny on the diversity and structure of insect-associated microbial communities was tested using two different approaches. First, Pagel’s λ (function *pgls()* within the package *caper)* and Blomberg’s *K* (function *phylosignal()* within the package *picante,* 9,999 permutations) were used to test whether bacterial diversity (Shannon index, averaged across replicated samples within the same host species) is phylogenetically conserved. Second, a Mantel test (9,999 permutations) was used to test the correlation between a Bray-Curtis matrix of the distance between insect host species calculated considering the composition of microbial communities (thus, averaged across replicated samples within the same host species) and a matrix of phylogenetic distance between insect species obtained using the *cophenetic.phylo*() from the *ape* R package (Paradis and Schliep, 2019).

The relative abundance of each bacterial genus *(n=* 18 for whole insects and *n=* 22 for guts) was fit using the *lmer()* function to test the effect of insect order, sex, life stage, sample origin, diet, and treatment. Data were merged according to the bacterial genus and filtered to remove those taxa that were representing < 1% of the whole dataset. Analyses were run separately for whole insects and insect gut, and also in this case study ID was included as a random factor (formula: *insect_order* + *sex* + *life_stage* + *diet* + *treatment* + *origin* + (1|*study_ID*)). The package *emmeans* was used to infer pairwise contrasts (corrected using False Discovery Rate).

## 3 Results

### 3.1 An overview of the studies included in the meta-analysis

Across the 246 studies included in this meta-analysis (and before filtering for the criteria indicated above) this survey identified 63 different DNA extraction methods, 44 primer pairs spanning all the 9 variable regions of the 16S rRNA, and 3 different sequencing technologies (Supplementary Materials 2). The majority of studies used the QIAGEN DNeasy Blood and Tissue Kit (*n*= 58), MOBIO PowerSoil kit (*n*= 28), and Phenol:Chloroform procedures (*n=* 23). The primer pair 515F-806R was the most widely used (*n*= 76), and amplicons were preferentially sequenced on Illumina technology (*n*= 181) using the MiSeq (*n*= 156) platform. Data were mostly submitted to SRA databases (*n*= 168). Most of the studies focused on single species (*n=* 142) rather than multiple species (*n*= 104).

On a negative note, several studies failed to report important information to ensure repeatability (e.g. DNA extraction method, primer pairs, sequencing platform, see Supplementary Material 2). This data was neither reported in the paper nor the SRA. Thus, the initial bulk of 246 studies was filtered to ensure high-quality standards for our dataset. First, studies were discarded if data were not publicly available, or if linked it was not available anyway (e.g. broken link) *(n*= 67, 27.24%), or not viable (i.e. non demultiplexed or not replicated, *n*= 16). To increase the comparability between studies in this meta-analysis, studies not focusing on the 16S rRNA (*n*= 2) and the V3-V4 region (*n*= 11), failing to report the primer pair they used (*n*= 16), or sequencing amplicons on platforms other than Illumina MiSeq (*n*= 47) were excluded. To ensure the easy repeatability of analysis, we also excluded those studies where data were made publicly available on a platform different than SRA (*n*= 14). This left a total of 72 studies to be included in the meta-analysis.

These 72 studies associate to 98 SRA BioProjects and, after QC on the sequencing data (see above), resulted in 5,212 unique samples. Most of the projects focused on the orders: Diptera (*n*= 15), Hymenoptera (*n*= 14), Coleoptera (*n*= 13), Hemiptera (*n*= 13) and Lepidoptera (*n*= 11), although the vast majority of samples belong to Hymenoptera (*n*= 1,525). Several projects focused on adult specimens (*n*= 60), collected in the field (*n*= 54), and specimen sex was often not reported (*n*= 61). Similarly, diet was often not reported (*n*= 29), or insects fed on plants (*n*= 32) or artificial diet (*n*= 21). Most projects focused on insect guts or gut’s sections (*n*= 52), but the majority of samples were obtained from whole insects (*n*= 2,442) or surface-sterilised insects (*n*= 1,872).

### 3.2 Which factors influence the structure and diversity of insect microbiota?

The analysis of the factors influencing the structure and the diversity of insect microbiota was performed on a subset of 2,442 samples for the group of whole insects and 1,872 samples for the group of insect guts, as explained above.

PERMANOVA analysis was used to test the amount of variance explained by each factor on the structure of insect microbiota using two different distance matrices to account for the relative abundance (Bray-Curtis distance matrix) or the presence/absence (Jaccard matrix). Results are very similar for both Bray-Curtis and Jaccard matrices, and whole insects and insect guts. While all considered factors (insect species, sex, life stage, diet, treatment, and origin) resulted to have a significant effect on the structure of insect microbiota (Tab. 1), insect species explained most of the variation (32.9-43.9%) while the other factors explained ≤ 1.5% (Tab. 1). The analysis of the effect of each factor on the Shannon diversity for each sample using a linear mixed-effects model showed that insect species was the factor still explaining most of the variation in the diversity of microbial communities in whole insects (35.1%) and insect guts (62.6%). Also, diet was an important factor explaining variation in bacterial community diversity, especially in guts. There was no effect of sample treatment on bacterial community diversity in whole insect specimens or guts.

**Table 1:** Analysis of the effect of insect species, sex, life stage, diet, sample treatment, and sample origin on the structure and diversity of insect-associated bacterial communities. The effect of each factor on the structure of microbial communities was tested using a PERMANOVA on a Bray-Curtis (accounting for relative abundance) or Jaccard (accounting for presence/absence) distance matrix. The effect of factors on microbial diversity was tested using a linear mixed-effects model on Shannon diversity index. Results are grouped by whole insect specimens (upper) or insect guts (lower).

| Factors (whole insect) | df | Bray-Curtis |  |  | Jaccard |  |  | Shannon |  |  |
| --- | --- | --- | --- | --- | --- | --- | --- | --- | --- | --- |
| | | $R^2$ | F | P | $R^2$ | F | P | $R^2$ | $\chi^2$ | P |
| Insect species | 134 | 0.428 | 13.2 | <0.001 | 0.335 | 8.8 | <0.001 | 0.351 | 1064.1 | <0.001 |
| Sex | 2 | 0.001 | 2.1 | <0.001 | 0.001 | 1.8 | <0.001 | 0.007 | 14.8 | <0.001 |
| Life stage | 4 | 0.006 | 6.5 | <0.001 | 0.005 | 4.8 | <0.001 | 0.038 | 40.4 | <0.001 |
| Diet | 4 | 0.008 | 8.5 | <0.001 | 0.006 | 5.8 | <0.001 | 0.034 | 27.2 | <0.001 |
| Treatment | 1 | 0.0004 | 1.9 | 0.01 | 0.0004 | 1.6 | 0.008 | 0.001 | 0.2 | 0.648 |
| Origin | 1 | 0.001 | 6.6 | <0.001 | 0.001 | 5.1 | <0.001 | 0.007 | 14.8 | <0.001 |
| Factors (guts) | df | $R^2$ | F | P | $R^2$ | F | P | $R^2$ | $\chi^2$ | P |
| Insect species | 89 | 0.439 | 16.5 | <0.001 | 0.329 | 10.1 | <0.001 | 0.626 | 343.9 | <0.001 |
| Sex | 2 | 0.003 | 5.5 | <0.001 | 0.002 | 4.05 | <0.001 | 0.001 | 4.8 | 0.089 |
| Life stage | 3 | 0.003 | 3.9 | <0.001 | 0.003 | 2.9 | <0.001 | 0.006 | 10.6 | 0.014 |
| Diet | 3 | 0.006 | 7.4 | <0.001 | 0.005 | 5.2 | <0.001 | 0.036 | 29.55 | <0.001 |
| Treatment | 1 | 0.001 | 3.2 | <0.001 | 0.001 | 2.7 | 0.008 | 0.0001 | 0.8 | 0.367 |
| Origin | 1 | 0.015 | 50.1 | <0.001 | 0.012 | 33.4 | <0.001 | 0.087 | 96.1 | <0.001 |

### 3.3 Is there a relationship between microbial diversity and insect phylogeny?

The results above suggest a strong influence of host species identity on the diversity and structure of insect-associated microbial communities, although the strength and direction of this effect remain unknown. This was further dissected by testing for a phylogenetic signal in the microbial community diversity (Shannon index) and structure (Bray-Curtis distance). This analysis just focused on insect guts (n=42) because the sample size for whole insects was not big enough to run any inference. A phylogeny of the included insect species is available as Supplementary Material 5. Results (λ = 0.735) suggest that bacterial diversity in insect guts is non-random (λ = 0, *P* < 0.001) but shows a weak phylogenetic signal (λ = 1, P < 0.001). These results were confirmed by the Blomberg’s *K* test (*K* = 0.17, P = 0.16). Similarly, Mantel’s test shows a weak correlation between the structure of microbiota and the host phylogenetic distance (*r* = 0.09, P = 0.015).

### 3.4 Response of bacterial taxa to different factors

To gain further insights on the variation of insect-associated bacterial communities, the factors insect order, sex, life stage, sample origin, diet, and treatment were tested against the relative abundance of each bacterial genus separately for whole insects and insect gut. Results are more extensively reported as Supplementary Material 6.

#### 3.4.1 Whole insects

When testing the effect of insect phylogenetic order on the relative abundance of the major bacterial genera identified in this meta-analysis, differences were found for the genus *Acinetobacter, Lactobacillus* and *Saccharibacter.* Specifically, *Lactobacillus* was more abundant in Hymenoptera when compared to Coleoptera, Diptera, Hemiptera, and Lepidoptera. While we observed a significant effect of insect order on the relative abundance of *Acinetobacter* (*P* = 0.02) and *Saccharibacter* (*P* = 0.02), the post-hoc analysis clarified that this was due to a marginal higher abundance of *Acinetobacter* in Coleoptera compared to Hemiptera (*P* = 0.05) and a marginal higher abundance of *Saccharibacter* in Hymenoptera when compared to Coleoptera (*P* = 0.06) and Diptera (*P* = 0.05).

The sex of the insect specimen influenced the relative abundance of *Arsenophonus, Salinibacter,* and *Wolbachia.* Although this observation was the result of differences between males/females towards samples with ‘unknown’ sex (see above), *Arsenophonus* resulted to be more abundant in males than females (*P* = 0.01). When testing the effect of life stage, *Enterococcus* showed a higher abudance in pupae when compared to juveniles (*P* < 0.001) and adults (*P* < 0.001), while it was found a higher abundance of *Leucobacter* in juveniles than any other stage (*P* ≤ 0.01) and of *Escherichia-Shigella* in juveniles compared to adults (*P* < 0.001).

When comparing samples from the field towards lab-reared insects, *Enterococcus* (*P* < 0.001) and *Pseudomonas* (*P* < 0.001) showed to be more abundant in insects grown in lab settings compared to those collected in the field. Diet was also an important factor that impacted several taxa. *Wolbachia* was more abundant in carnivores and insects feeding on artificial diet than herbivores and those feeding on fungi (*P* < 0.001). *Pseudomonas* was more abundant in insect feeding on fungi or artificial diet than carnivores and herbivores (*P* ≤ 0.02), while the opposite was observed for *Escherichia-Shigella* (*P* < 0.001). *Buchnera* was more abundant on carnivores (*P* < 0.001) while *Enterococcus* was least abundant on carnivores compared to all the other groups (*P* ≤ 0.03). When considering sample treatment as a factor, *Salinibacter* (*P* = 0.002) was more abundant in samples subjected to treatment while *Leucobacter* (*P* = 0.01) was more abundant in control samples.

#### 3.4.2 Insect guts

When focusing only on insect guts, *Alistipes, Bacteroides, Desulfovibrio, Christensenellaceae R-7 group* were consistently more abundant (*P* < 0.05) in Blattodea than Coleoptera, Diptera, Hemiptera, Hymenoptera, and Lepidoptera. On the other hand, *Providencia* was consistently more abundant (*P* < 0.01) in Diptera than Coleoptera, Blattodea, Hemiptera, Hymenoptera, and Lepidoptera. Similarly, *Snodgrassella* was more abundant in Hymenoptera (*P* < 0.05) than Coleoptera, Hemiptera, Hymenoptera, and Lepidoptera. The significant effect of insect order on *Lactobacillus* was probably driven from a marginal higher abundance of this genus in Hymenoptera compared to Coleoptera (*P* = 0.08), Diptera (*P* = 0.05), and Lepidoptera (*P* = 0.06). Similarly, *Gilliamella* was more abundant in n Hymenoptera compared to Coleoptera (*P* = 0.06) and Diptera (*P* = 0.05). *Wolbachia* was marginally more abundant in Hemiptera than Coleoptera and Diptera (*P* = 0.05).

Contrasting male and female specimens, *Lactobacillus* (*P* = 0.006) and *Blattabacterium* (*P* = 0.006) were more abundant in females, while *Enterobacter* (*P* = 0.02) and *Christensenellaceae R-7 group* (*P* = 0.009) were more abundant in male specimens. Comparing life stages, results suggest a consistent lower abundance of *Providencia* in eggs compared to all other instars (*P* < 0.01), and in pupae when compared to juveniles (*P* = 0.01). *Pseudomonas* was consistently more abundant in juveniles compared to adults (*P* = 0.01). *Acinetobacter* and *Enterobacter* were more abundant in adults than juveniles (*P* < 0.001), while *Klebsiella* was more abundant in juveniles compared to adults and pupae (*P* < 0.001).

The comparison between samples collected in the field and specimens coming from laboratory cultures highlighted that *Lactobacillus*, *Enterococcus*, *Providencia*, and *Blattabacterium* were more abundant in specimens from the laboratory (*P* ≤ 0.05), while *Alistipes, Acinetobacter, Desulfovibrio, Bacteroides,* and *Christensenellaceae R-7 group* were more abundant in specimens from the field (*P* ≤ 0.003). When focusing on the diet, results suggest that *Providencia* was more abundant in insects fed with artificial diet than herbivores or carnivores (*P* < 0.001), and *Wolbachia* was more abundant in carnivores than herbivores (*P* < 0.001). Several other groups did show differential abundance according to the diet, but the post-hoc contrasts highlighted differences towards a group of samples which diet was not specified in the study. *Snodgrassella* was more abundant (*P* < 0.001) in control groups than treated insects, while those subjected to treatment showed an increase of *Acinetobacter* (*P* < 0.004) and *Gilliamella* (*P* < 0.004).

## 4 Discussion

This is the first study that provides a comprehensive analysis of insect-associated bacterial communities by screening 246 studies and analysing >4,000 samples under the same framework. This analysis reveals a wide range of approaches that have been used to study insect microbiota, which leads to issues when attempting a comparison across studies. Also, insect species was identified as the factor most impacting the structure and diversity of insect bacterial communities. However, this study finds weak support for insect-microbiota co-diversification.

### 4.1 Methodological considerations

This meta-analysis screened 246 studies that focused on the analysis of insect-associated bacterial communities using 16S-amplicon HTS, identifying several DNA extraction techniques (63) and primer pairs (44).

DNA extraction has been identified as one of the factors that influence the outcome of metabarcoding analyses (McOrist et al., 2002; Walker et al., 2015; Vasselon et al., 2017; Hallmaier-Wacker et al., 2018; Dopheide et al., 2019; Maillet et al., 2021, but see Fouhy et al. 2016). The fact that different methods have a different impact on the final reconstruction of the microbial community might reside on the efficiency of DNA recovery or on the ability of different protocols to isolate the DNA from cells of specific groups (e.g., gram+ bacteria). Also, different DNA extraction procedures might perform differently in purifying nucleic acids from PCR inhibitors, which can be an important factor influencing the quality of the amplicon library. Unfortunately, there is no accurate study that links specific DNA extraction methods to the presence/absence of specific bacterial groups, thus we are still not able to predict which taxa might be over-/under-represented when using different protocols.

While standardizing the DNA extraction method for entire fields represents a big challenge, it should be definitely proposed for PCR primer pairs. The influence of primer pairs on the outcome of metabarcoding analyses is not new to the literature (Tremblay et al., 2015; Parada et al., 2016; Bahram et al., 2019). A recent study compared the efficiency of primer pairs amplifying different regions of the bacterial 16S rRNA on the correct reconstruction of complex microbial communities (Abellan-Schneyder et al., 2021). They found differences in the performance of different primer pairs in reconstructing mock communities, with some of them failing to amplify entire microbial groups. Thus, there is an increasing need to run preliminary tests of different primer combinations when working with new systems, while this limitation might be overcome by sequencing the full length of the 16S rRNA (e.g. using PacBio or Oxford Nanopore sequencing technologies).

In addition, this study revealed even more concerning aspects. About 30% of the considered studies did not publicly share the raw datasets, or totally failed to report important information that would ensure repeatability and comparability (e.g. DNA extraction method, amplified region, sequencing platform). Whilst the benefits of Public Data Archiving are clear and widely acknowledged, we are still very far from seeing it as a common practice (Parr and Cummings, 2005; Roche et al., 2015; Whitlock, 2011).

A caveat of this study is the limited power to correct the data for possible contaminations, which may occur via instrument/reagents contamination and index hopping. Unfortunately, a careful account of contamination was not possible because just a few studies included non-template control samples, and discarding those that do not include such samples in their design would completely hinder the feasibility of this work. However, the workflow included a series of steps that helped in accounting for possible contaminations, regardless their source. First, “study ID” was included as random effect in most of the analyses, and this helped to correct the results for study-specific effects, which also include study-specific contaminations. Second, this analysis builds on a massive amount of data ( >4,000 samples), and studies/samples were carefully selected to ensure that all the groups had a sample size guaranteeing a solid statistical inference. Having such sample size helped in counteracting contamination effects that might have occurred for some studies/samples. Third, before testing the influence of different factors on microbial taxa, those bacterial genera that represented <1% of the whole community for each sample were discarded, further helping to remove the effect of minor contaminations and index hopping, if any. Fourth, this meta-analysis acknowledges the limits of metabarcoding and – as a precaution – analyses and discussions on bacterial taxa were not pushed beyond the genus level, limiting the impact of possible contaminations on the overall findings of this study.

### 4.2 Factors influencing insect microbial communities

This study suggests that insect species are the major factor that shapes the structure and diversity of insect-associated bacterial communities. The analysis considered also diet, sex, life stage, treatment (control or treated group, regardless of the treatment), and origin (lab or field sample) as factors, and all of them had a significant effect on the structure and diversity of bacterial communities, except treatment which did not affect the diversity of insect bacterial communities. However, looking at the variance explained by each factor, it appears clearly that insect species represent the most important factor in structuring insect microbial communities. This is followed by diet, although at a much lower magnitude. This result was consistent in both whole insects and insect guts. While all the factors considered had a significant effect, this might be driven by the very high sample size (>4,000 samples) but with a scarce or context-dependent functional meaning. This hypothesis is corroborated by the results obtained when analyzing the data using the DADA2 pipeline (Supplementary Material 4). In this case, the sample size was lower (~ 2,000 samples) due to more strict QC, and indeed results suggest that some factors do not significantly influence the diversity/structure of insect microbiota (e.g. treatment), although the results still hold for those explaining a higher portion of the variance (e.g. insect species, diet) which are more likely to have a functional meaning.

Several previous studies tested whether different insect species (mostly within the same genus or trophic niche) associate with different microbial communities. Differences between insect species have been found in aphids (Zepeda-Paulo et al., 2018; Fakhour et al., 2018; McLean et al., 2019; Gallo-Franco et al., 2019), mealybugs (Lin et al., 2019), reduviids (Rodríguez-Ruano et al., 2018), fruit flies (Ventura et al., 2018; Yong et al., 2017), beetles (Hulcr et al., 2012; Kolasa et al., 2019), silkworms (Chen et al., 2018), Lepidopera (van Schooten et al., 2018; Liu et al., 2020), mosquitoes (Mancini et al., 2018), bees and wasps (Skrodenytė-Arbačiauskienė et al., 2019; Suenami et al., 2019). Conversely, other studies did not report differences when comparing different insect species (Berasategui et al., 2016; Ramalho et al., 2017; Meng et al., 2019a). Together with the result that host species explains most of the variability of insect-associate microbial communities, this might suggest the existence of a relationship between host phylogeny and the diversity/structure of insect-associated microbiota.

If this would be the case, we should be able to observe a signal of phylosymbiosis – the relationship between the host phylogeny and the structure of their microbial communities (Mazel et al., 2018; Lim and Bordenstein, 2020). However, when specifically testing for signatures of phylosymbiosis, results revealed a weak phylogenetic signal on the diversity and structure of insect-associated microbiota. This is not surprising, as similar results have been found in several other lineages (Mazel et al., 2018; Chiarello et al., 2018; Lutz et al., 2019; Grond et al., 2020; Trevelline et al., 2020). The fact that host species identity is the most important factor in structuring insect-associated microbial communities, while there is weak support for phylosymbiosis, might suggest that each insect species represents a unique combination of traits that support a unique microbiota. This might have ecological consequences, as a microbiome that is not tied to host phylogeny might also be important to colonize new ecological niches. This idea, although needs to be tested more accurately, might fit well with the diversity of insects. Indeed, insects are pretty unique in this sense, as their morphology, diet, niche, and behavior greatly varies between closely related species (e.g. specialist vs. generalist fruit flies) or even within the same species across development (e.g. dragonflies, hoverflies) or between sexes (e.g. mosquitoes). This high intra- and interspecific variability can ultimately result in weak or no phylosymbiosis. However, signals of phylosymbiosis have been found when focusing on single insect genera, for example, *Nasonia, Drosophila,* and few mosquitoes lineages (Brooks et al., 2016). Thus, it is possible that phylosymbiosis cannot be detected when working at a large phylogenetic scale, but rather in specific sub-clades. Indeed, the patterns of microbiome acquisition greatly vary across clades, and there is evidence that it can even be acquired from the environment (Rassati et al., 2019). An alternative hypothesis might be that while phylosymbiosis does not broadly happen in insects, perhaps the functionality of the microbiomes (i.e., gene content) actually reconciles with the host phylogeny. Indeed, previous studies found that an organism can interact with phylogenetically distinct microorganisms that provide similar core functions (Parfrey et al., 2018; Roth-Schulze et al., 2018). This idea needs to be further tested with the analyses enabled by meta’omics techniques (Malacrinò, 2018).

### 4.3 Response of bacterial clades to factors influencing insect microbial communities

While insect species was the strongest driver of structure and diversity of bacterial communities in this study, diet also showed a significative effect. Several previous studies showed that insect-associated microbial communities are often shaped by their host’s diet, for example in termites (Mikaelyan et al., 2015), aphids (Wu et al., 2018; Xu et al., 2019, 2020; Holt et al., 2020), bugs (Martinez et al., 2019), beetles (Kim et al., 2017), fruit flies (Malacrinò et al., 2018; Majumder et al., 2019; Woruba et al., 2019; Asimakis et al., 2019), soldier flies (Bruno et al., 2019) and ants (Russell et al., 2009). Interestingly, the bacterial communities associated with *Phtorimaea operculella* varied even according to the plant tissue insects were feeding on (Zheng et al., 2020). In *Agrilus mali,* Zhang et al. (2018b) found differences in the insect-associated fungal community according to the host plant, but not in the bacterial community. Diet-dependent microbiota has been reported for several species of Lepidoptera although, recently, it has been suggested that Lepidoptera lack a resident microbiota, and their gut microbiota resembles the one of the host plants or is assembled from the environmental microbiota (Montagna et al., 2016; Whitaker et al., 2016; Staudacher et al., 2016; Hammer et al., 2017; Phalnikar et al., 2018; Strano et al., 2018; Minard et al., 2019; Liu et al., 2020). Furthermore, there is evidence showing that the bacterial microbiota of cockroaches is very stable and is not influenced by diet (Tinker and Ottesen, 2016; Lampert et al., 2019, but see Pérez-Cobas et al. 2015). Similarly, no diet-mediated effect has been reported in houseflies microbiota (Xue et al., 2019). Results suggest a higher abundance of *Wolbachia* and *Buchnera* in carnivores compared to other groups. This effect might be driven by the diet or, in alternative, this can be explained by the fact that all carnivores in this meta-analysis are hemipterans, which is an order where *Wolbachia* and *Buchnera* represent major symbionts. The higher abundance of *Pseudomonas* on insects feeding on an artificial diet might be due to a dysbiosis driven by not feeding on their natural substrate, but further tests are needed to test this hypothesis.

The life stage represented another factor that influences the structure and diversity of insect microbial communities. Johnston and Rolff (2015) suggest that both the insect and its symbionts modulate the gut microbiota during development. A very wide variety of studies confirms that insect-associated bacterial community varies across developmental species in dragonflies (Nobles and Jackson, 2020), cockroaches (Carrasco et al., 2014), thrips (Kaczmarczyk et al., 2018), lacewings (Zhao et al., 2019), psyllids (Meng et al., 2019b), beetles (Briones-Roblero et al., 2017; Kim et al., 2017; Wang et al., 2018; Wilches et al., 2018; Mason et al., 2019; Chouaia et al., 2019; Ali et al., 2019), fruit flies (Malacrinò et al., 2018; Andongma et al., 2019; Yao et al., 2019; Huang et al., 2019; Gallo-Franco and Toro-Perea, 2020), mosquitoes (Pennington et al., 2016), and moths (Chen et al., 2016; González-Serrano et al., 2019). While there are some reports of changes of insect bacterial communities across life stages in Lepidoptera, Phalnikar et al. (2018) investigating twelve butterfly species found little changes in some species and wider changes in others, with the wider impact driven by insect species. Similarly, very limited changes in bacterial microbiota have been found across different stages of *Plodia interpunctella* (Mereghetti et al., 2019) and blow flies (Wohlfahrt et al., 2020). The higher abundance of *Enterococcus* in pupae observed in this meta-analysis might enhance the defenses against pathogens during this vulnerable developmental stage (Johnston and Rolff, 2015; Grau et al., 2017).

The wider literature on host-microbiota interactions often reports that microbial communities associated with any other organisms are subjected to shifts when an external input is applied. These inputs can be physical (e.g. temperature), chemical (e.g. xenobiotics), or biological (e.g. microorganisms), and they influence both the host and its microbiota. This is also the case in insects, where literature suggests an effect of antibiotics (Hammer et al., 2016; Guégan et al., 2018; Koskinioti et al., 2019), agrochemicals (Kakumanu et al., 2016; Raymann et al., 2018; Receveur et al., 2018; Akami et al., 2019; Botina et al., 2019; Ding et al., 2019; Zhang and Yang, 2019; Li et al., 2020a), heavy metals (Zhu et al., 2018b; Rothman et al., 2019a,b; Li et al., 2020b), land use (Zhu et al., 2018a), pollutants (Duguma et al., 2017; Cassone et al., 2020), other microorganisms (Billiet et al., 2017; Sun et al., 2018; Zhang et al., 2018a; Vacheron et al., 2019) and irradiation (Cai et al., 2018; Stathopoulou et al., 2019) on insect-associated microbial communities. Interestingly, this study shows differences between control and treatment (regardless of the nature of the treatment) in the structure but not in the diversity of insect-associated microbial communities. A possible explanation for this effect is that treatments disrupt the equilibrium in the insect microbiota, resulting in shifts in the community structure but not in the diversity of the members that create the community. Unfortunately, the current knowledge does not allow to speculate about the changes of *Salinibacter* and *Leucobacter* as an effect of treatment.

While sex is often not considered when studying insect microbial communities, there is evidence of differences in microbiota composition between females and males in *Leptocybe invasa* (Guo et al., 2020), *Dendroctonus valens* (Xu et al., 2016), *Anopheles gambiae,* and *Anopheles coluzzii* (Segata et al., 2016). It is not surprising that *Arsenophonus* resulted to be differentially abundant between males and females, as species of this bacterial clade are known for their ability to manipulate sex ratios in various insect species (Kageyama et al., 2012). Also, few studies tested whether rearing insects in laboratory settings can have an influence on their microbial communities, and there is the suggestion that such differences exist (Hegde et al., 2018; Kakumanu et al., 2018; Hadapad et al., 2019; Waltmann et al., 2019; Gomes et al., 2020). Very likely these differences are mainly a consequence of a diet effect, given that many insects are grown on an artificial diet in the laboratory, as suggested by a higher abundance of *Enterococcus* and *Pseudomonas* in laboratory-reared samples overlapping the results of insects feeding on artificial diet.

### 5 Conclusions

This study shows that host species is the most important driver of the structure and diversity of insect-associated bacterial communities. While confirming previous reports (Colman et al., 2012; Jones et al., 2013; Yun et al., 2014), this work expands by testing the effects of a wider set of factors on the insect-associated microbial assemblies, suggesting that their magnitude is marginal compared to the host species-driven effect.

Importantly, this study found several gaps that need to be considered in future research endeavours. Indeed, there is very small consistency in several experimental procedures (e.g. DNA extraction, PCR primers choice) that might lead to bias, artifacts and, in particular, it might prevent the comparison between different experiments. Also, the raw data were often improperly compiled or not shared at all. Besides the impact on comparability and repeatability, this has a huge cost. Data that are properly collected and shared speeds up scientific research (i.e., no new experiments needed), saves taxpayers’ money (i.e., no experiment is needed if data to test that hypothesis already exist), has a lower environmental impact (i.e., fewer experiments equals fewer consumables to dispose of) and reduces inequality (i.e., allows to test hypotheses when there is no sufficient funding to perform experiments).

Furthermore, there is an evident bias towards insect species and orders with economical importance, and this leads to a low representation of much of the insect diversity. There is much need to investigate the microbial diversity of insects belonging to different orders to increase our ability to understand microbial community assembly. Also, most of the papers focused on the bacterial community and, as often happened in other fields, other communities (e.g. fungal) are not investigated although we are aware of their major role in insect biology and fitness.

The fact the insect species is the most important driver of insect microbiota, followed by a wide range of other factors, increases our understanding of the biology of this clade of organisms so important for a variety of ecosystems. Also, a deep understanding of the assembly and function of these microbial communities can have a positive impact on insect conservation, pest management, and biosurveillance (Malacrinò, 2018; Trevelline et al., 2019). This study also witnesses the efforts of the scientific community in understanding the taxonomical structure and diversity of insect-associated microbial communities. While this approach is powerful and helped to gain a wide understanding of the dynamics of insect-associated microbes, it does not tell us the functional role of those microbial communities leaving the question *‘what do they do?’* still unanswered. Thus, it is timely to push our endeavours towards a deeper understanding of the microbial gene content and expression using the variety of meta’omics tools currently available.

## Data availability

Raw data from single studies are publicly available. The code used to perform analyses has been deposited on GitHub: https://github.com/amalacrino/insect_microbiota_metaanalysis

## Conflict of interests

I declare that no competing interests exist.

## Supplementary materials

**Supplementary material 1**. List of the studies included in this meta-analysis.

**Supplementary material 2**. Summary of the studies included in this meta-analysis.

**Supplementary material 3**. Summary of the samples analysed in this meta-analysis.

**Supplementary material 4**. Results obtained from running the analysis after processing the raw data using the DADA2 pipeline.

**Supplementary material 5**. Phylogenetic tree showing the range of insect species used in testing whether there is a relationship between microbial diversity and insect phylogeny.

**Supplementary material 6**. Results from testing the influence of different factors (insect order, diet, sex, life stage, treatment, and origin) on the relative abundance of bacterial genera.

## Notes

https://github.com/amalacrino/insect_microbiota_metaanalysis

### Competing Interest Statement

The authors have declared no competing interest.

## References

Abellan-Schneyder, I., Matchado, M. S., Reitmeier, S., Sommer, A., Sewald, Z., Baumbach, J., List, M., and Neuhaus, K. (2021). Primer, pipelines, parameters: Issues in 16s rrna gene sequencing. mSphere, 6(1).

Akami, M., Njintang, N. Y., Gbaye, O. A., Andongma, A. A., Rashid, M. A., Niu, C.-Y., and Nukenine, E. N. (2019). Gut bacteria of the cowpea beetle mediate its resistance to dichlorvos and susceptibility to lippia adoensis essential oil. Scientific reports, 9(1):1–13.

Ali, H., Muhammad, A., Sanda, N. B., Huang, Y., and Hou, Y. (2019). Pyrosequencing uncovers a shift in bacterial communities across life stages of octodonta nipae (coleoptera: Chrysomelidae). Frontiers in microbiology, 10:466.

Andongma, A. A., Wan, L., Dong, Y.-C., Wang, Y.-L., He, J., and Niu, C.-Y. (2019). Assessment of the bacteria community structure across life stages of the chinese citrus fly, bactrocera minax (diptera: Tephritidae). BMC microbiology, 19(1):1–9.

Asimakis, E. D., Khan, M., Stathopoulou, P., Caceres, C., Bourtzis, K., and Tsiamis, G. (2019). The effect of diet and radiation on the bacterial symbiome of the melon fly, zeugodacus cucurbitae (coquillett). BMC biotechnology, 19(2):1–12.

Bahram, M., Anslan, S., Hildebrand, F., Bork, P., and Tedersoo, L. (2019). Newly designed 16s rrna metabarcoding primers amplify diverse and novel archaeal taxa from the environment. Environmental Microbiology Reports, 11(4):487–494.

Bartoń, K. (2009). MuMIn: Multi-model inference. R package version 0.12.2.

Bates, D., Mächler, M., Bolker, B., and Walker, S. (2014). Fitting linear mixed-effects models using lme4. arXiv preprint arXiv:1406.5823.

Berasategui, A., Axelsson, K., Nordlander, G., Schmidt, A., Borg-Karlson, A.-K., Gershenzon, J., Terenius, O., and Kaltenpoth, M. (2016). The gut microbiota of the pine weevil is similar across europe and resembles that of other conifer-feeding beetles. Molecular ecology, 25(16):4014–4031.

Billiet, A., Meeus, I., Cnockaert, M., Vandamme, P., Van Oystaeyen, A., Wäckers, F., and Smagghe, G. (2017). Effect of oral administration of lactic acid bacteria on colony performance and gut microbiota in indoor-reared bumblebees (bombus terrestris). Apidologie, 48(1):41–50.

Botina, L., Vélez, M., Barbosa, W., Mendonça, A., Pylro, V., Tótola, M., and Martins, G. (2019). Behavior and gut bacteria of partamona helleri under sublethal exposure to a bioinsecticide and a leaf fertilizer. Chemosphere, 234:187–195.

Briones-Roblero, C. I., Hernández-García, J. A., Gonzalez-Escobedo, R., Soto-Robles, L. V., Rivera-Orduña, F. N., and Zúñiga, G. (2017). Structure and dynamics of the gut bacterial microbiota of the bark beetle, dendroctonus rhizophagus (curculionidae: Scolytinae) across their life stages. PloS one, 12(4):e0175470.

Brooks, A. W., Kohl, K. D., Brucker, R. M., van Opstal, E. J., and Bordenstein, S. R. (2016). Phylosymbiosis: relationships and functional effects of microbial communities across host evolutionary history. PLoS biology, 14(11):e2000225.

Bruno, D., Bonelli, M., De Filippis, F., Di Lelio, I., Tettamanti, G., Casartelli, M., Ercolini, D., and Caccia, S. (2019). The intestinal microbiota of hermetia illucens larvae is affected by diet and shows a diverse composition in the different midgut regions. Applied and environmental microbiology, 85(2).

Cai, Z., Yao, Z., Li, Y., Xi, Z., Bourtzis, K., Zhao, Z., Bai, S., and Zhang, H. (2018). Intestinal probiotics restore the ecological fitness decline of bactrocera dorsalis by irradiation. Evolutionary applications, 11(10):1946–1963.

Callahan, B. J., McMurdie, P. J., Rosen, M. J., Han, A. W., Johnson, A. J. A., and Holmes, S. P. (2016). Dada2: high-resolution sample inference from illumina amplicon data. Nature methods, 13(7):581–583.

Carrasco, P., Pérez-Cobas, A. E., Van de Pol, C., Baixeras, J., Moya, A., and Latorre, A. (2014). Succession of the gut microbiota in the cockroach blattella germanica. Int Microbiol, 17(2):99–109.

Cassone, B. J., Grove, H. C., Elebute, O., Villanueva, S. M., and LeMoine, C. M. (2020). Role of the intestinal microbiome in low-density polyethylene degradation by caterpillar larvae of the greater wax moth, galleria mellonella. Proceedings of the Royal Society B, 287(1922):20200112.

Chen, B., Du, K., Sun, C., Vimalanathan, A., Liang, X., Li, Y., Wang, B., Lu, X., Li, L., and Shao, Y. (2018). Gut bacterial and fungal communities of the domesticated silkworm (bombyx mori) and wild mulberry-feeding relatives. The ISME journal, 12(9):2252–2262.

Chen, B., Teh, B.-S., Sun, C., Hu, S., Lu, X., Boland, W., and Shao, Y. (2016). Biodiversity and activity of the gut microbiota across the life history of the insect herbivore spodoptera littoralis. Scientific reports, 6(1):1–14.

Chiarello, M., Auguet, J.-C., Bettarel, Y., Bouvier, C., Claverie, T., Graham, N. A., Rieuvilleneuve, F., Sucré, E., Bouvier, T., and Villéger, S. (2018). Skin microbiome of coral reef fish is highly variable and driven by host phylogeny and diet. Microbiome, 6(1):1–14.

Chouaia, B., Goda, N., Mazza, G., Alali, S., Florian, F., Gionechetti, F., Callegari, M., Gonella, E., Magoga, G., Fusi, M., et al. (2019). Developmental stages and gut microenvironments influence gut microbiota dynamics in the invasive beetle popillia japonica newman (coleoptera: Scarabaeidae). Environmental microbiology, 21(11):4343–4359.

Colman, D. R., Toolson, E. C., and Takacs-Vesbach, C. (2012). Do diet and taxonomy influence insect gut bacterial communities? Molecular ecology, 21(20):5124–5137.

Dillon, R. J. and Dillon, V. (2004). The gut bacteria of insects: nonpathogenic interactions. Annual Reviews in Entomology, 49(1):71–92.

Ding, J., Zhu, D., Chen, Q.-L., Zheng, F., Wang, H.-T., and Zhu, Y.-G. (2019). Effects of longterm fertilization on the associated microbiota of soil collembolan. Soil Biology and Biochemistry, 130:141–149.

Dixon, P. (2003). Vegan, a package of r functions for community ecology. Journal of Vegetation Science, 14(6):927–930.

Dopheide, A., Xie, D., Buckley, T. R., Drummond, A. J., and Newcomb, R. D. (2019). Impacts of dna extraction and pcr on dna metabarcoding estimates of soil biodiversity. Methods in Ecology and Evolution, 10(1):120–133.

Douglas, A. E. (2009). The microbial dimension in insect nutritional ecology. Functional Ecology, 23:38–47.

Douglas, A. E. (2015). Multiorganismal insects: diversity and function of resident microorganisms. Annual review of entomology, 60:17.

Duguma, D., Hall, M. W., Smartt, C. T., and Neufeld, J. D. (2017). Effects of organic amendments on microbiota associated with the culex nigripalpus mosquito vector of the saint louis encephalitis and west nile viruses. Msphere, 2(1).

Fakhour, S., Ambroise, J., Renoz, F., Foray, V., Gala, J.-L., and Hance, T. (2018). A large-scale field study of bacterial communities in cereal aphid populations across morocco. FEMS microbiology ecology, 94(3):fiy003.

Fouhy, F., Clooney, A. G., Stanton, C., Claesson, M. J., and Cotter, P. D. (2016). 16s rrna gene sequencing of mock microbial populations-impact of dna extraction method, primer choice and sequencing platform. BMC microbiology, 16(1):1–13.

Gallo-Franco, J. J., Duque-Gamboa, D. N., and Toro-Perea, N. (2019). Bacterial communities of aphis gossypii and myzus persicae (hemiptera: Aphididae) from pepper crops (capsicum sp.). Scientific reports, 9(1):1–12.

Gallo-Franco, J. J. and Toro-Perea, N. (2020). Variations in the bacterial communities in anastrepha obliqua (diptera: Tephritidae) according to the insect life stage and host plant. Current microbiology, 77(7):1283–1291.

Gomes, A. F. F., Omoto, C., and Cônsoli, F. L. (2020). Gut bacteria of field-collected larvae of spodoptera frugiperda undergo selection and are more diverse and active in metabolizing multiple insecticides than laboratory-selected resistant strains. Journal of Pest Science, 93(2):833–851.

González-Serrano, F., Pérez-Cobas, A. E., Rosas, T., Baixeras, J., Latorre, A., and Moya, A. (2019). The gut microbiota composition of the moth brithys crini reflects insect metamorphosis. Microbial ecology, pages 1–11.

Grau, T., Vilcinskas, A., and Joop, G. (2017). Probiotic enterococcus mundtii isolate protects the model insect tribolium castaneum against bacillus thuringiensis. Frontiers in microbiology, 8:1261.

Grond, K., Bell, K. C., Demboski, J. R., Santos, M., Sullivan, J. M., and Hird, S. M. (2020). No evidence for phylosymbiosis in western chipmunk species. FEMS microbiology ecology, 96(1):fiz182.

Guégan, M., Minard, G., Tran, F.-H., Tran Van, V., Dubost, A., and Valiente Moro, C. (2018). Short-term impacts of anthropogenic stressors on aedes albopictus mosquito vector microbiota. FEMS microbiology ecology, 94(12):fiy188.

Guo, C., Peng, X., Zheng, X., Wang, X., Wang, R., Huang, Z., and Yang, Z. (2020). Comparison of bacterial diversity and abundance between sexes of leptocybe invasa fisher & la salle (hymenoptera: Eulophidae) from china. PeerJ, 8:e8411.

Gurung, K., Wertheim, B., and Falcao Salles, J. (2019). The microbiome of pest insects: it is not just bacteria. Entomologia Experimentalis et Applicata, 167(3):156–170.

Hadapad, A. B., Shettigar, S. K., and Hire, R. S. (2019). Bacterial communities in the gut of wild and mass-reared zeugodacus cucurbitae and bactrocera dorsalis revealed by metagenomic sequencing. BMC microbiology, 19(1):1–11.

Hallmaier-Wacker, L. K., Lueert, S., Roos, C., and Knauf, S. (2018). The impact of storage buffer, dna extraction method, and polymerase on microbial analysis. Scientific reports, 8(1):1–9.

Hammer, T. J., Fierer, N., Hardwick, B., Simojoki, A., Slade, E., Taponen, J., Viljanen, H., and Roslin, T. (2016). Treating cattle with antibiotics affects greenhouse gas emissions, and microbiota in dung and dung beetles. Proceedings of the Royal Society B: Biological Sciences, 283(1831):20160150.

Hammer, T. J., Janzen, D. H., Hallwachs, W., Jaffe, S. P., and Fierer, N. (2017). Caterpillars lack a resident gut microbiome. Proceedings of the National Academy of Sciences, 114(36):9641–9646.

Hegde, S., Khanipov, K., Albayrak, L., Golovko, G., Pimenova, M., Saldaña, M. A., Rojas, M. M., Hornett, E. A., Motl, G. C., Fredregill, C. L., et al. (2018). Microbiome interaction networks and community structure from laboratory-reared and field-collected aedes aegypti, aedes albopictus, and culex quinquefasciatus mosquito vectors. Frontiers in microbiology, 9:2160.

Holt, J. R., Styer, A., White, J. A., Armstrong, J. S., Nibouche, S., Costet, L., Malacrinò, A., Antwi, J. B., Wulff, J., Peterson, G., et al. (2020). Differences in microbiota between two multilocus lineages of the sugarcane aphid (melanaphis sacchari) in the continental united states. Annals of the Entomological Society of America, 113(4):257–265.

Huang, H., Li, H., Ren, L., and Cheng, D. (2019). Microbial communities in different developmental stages of the oriental fruit fly, bactrocera dorsalis, are associated with differentially expressed peptidoglycan recognition protein-encoding genes. Applied and environmental microbiology, 85(13).

Hulcr, J., Rountree, N., Diamond, S., Stelinski, L., Fierer, N., and Dunn, R. (2012). Mycangia of ambrosia beetles host communities of bacteria. Microbial ecology, 64(3):784–793.

Johnston, P. R. and Rolff, J. (2015). Host and symbiont jointly control gut microbiota during complete metamorphosis. PLoS Pathog, 11(11):e1005246.

Jones, R. T., Sanchez, L. G., and Fierer, N. (2013). A cross-taxon analysis of insect-associated bacterial diversity. PLoS one, 8(4):e61218.

Kaczmarczyk, A., Kucharczyk, H., Kucharczyk, M., Kapusta, P., Sell, J., and Zielińska, S. (2018). First insight into microbiome profile of fungivorous thrips hoplothrips carpathicus (insecta: Thysanoptera) at different developmental stages: molecular evidence of wolbachia endosymbiosis. Scientific reports, 8(1):1–13.

Kageyama, D., Narita, S., and Watanabe, M. (2012). Insect sex determination manipulated by their endosymbionts: incidences, mechanisms and implications. Insects, 3(1):161–199.

Kakumanu, M. L., Maritz, J. M., Carlton, J. M., and Schal, C. (2018). Overlapping community compositions of gut and fecal microbiomes in lab-reared and field-collected german cockroaches. Applied and Environmental Microbiology, 84(17).

Kakumanu, M. L., Reeves, A. M., Anderson, T. D., Rodrigues, R. R., and Williams, M. A. (2016). Honey bee gut microbiome is altered by in-hive pesticide exposures. Frontiers in microbiology, 7:1255.

Kim, J. M., Choi, M.-Y., Kim, J.-W., Lee, S. A., Ahn, J.-H., Song, J., Kim, S.-H., and Weon, H.-Y. (2017). Effects of diet type, developmental stage, and gut compartment in the gut bacterial communities of two cerambycidae species (coleoptera). Journal of Microbiology, 55(1):21–30.

Kolasa, M., Ścibior, R., Mazur, M. A., Kubisz, D., Dudek, K., and Kajtoch, Ł. (2019). How hosts taxonomy, trophy, and endosymbionts shape microbiome diversity in beetles. Microbial ecology, 78(4):995–1013.

Koskinioti, P., Ras, E., Augustinos, A. A., Tsiamis, G., Beukeboom, L. W., Caceres, C., and Bourtzis, K. (2019). The effects of geographic origin and antibiotic treatment on the gut symbiotic communities of bactrocera oleae populations. Entomologia Experimentalis et Applicata, 167(3):197–208.

Kumar, S., Stecher, G., Suleski, M., and Hedges, S. B. (2017). Timetree: a resource for timelines, timetrees, and divergence times. Molecular biology and evolution, 34(7):1812–1819.

Lampert, N., Mikaelyan, A., and Brune, A. (2019). Diet is not the primary driver of bacterial community structure in the gut of litter-feeding cockroaches. BMC microbiology, 19(1):1–14.

Lewis, Z. and Lizé, A. (2015). Insect behaviour and the microbiome. Current Opinion in Insect Science, 9:86–90.

Li, F., Li, M., Mao, T., Wang, H., Chen, J., Lu, Z., Qu, J., Fang, Y., Gu, Z., and Li, B. (2020a). Effects of phoxim exposure on gut microbial composition in the silkworm, bombyx mori. Ecotoxicology and environmental safety, 189:110011.

Li, M., Li, F., Lu, Z., Fang, Y., Qu, J., Mao, T., Wang, H., Chen, J., and Li, B. (2020b). Effects of tio2 nanoparticles on intestinal microbial composition of silkworm, bombyx mori. Science of the Total Environment, 704:135273.

Lim, S. J. and Bordenstein, S. R. (2020). An introduction to phylosymbiosis. Proceedings of the Royal Society B, 287(1922):20192900.

Lin, D., Zhang, L., Shao, W., Li, X., Liu, X., Wu, H., and Rao, Q. (2019). Phylogenetic analyses and characteristics of the microbiomes from five mealybugs (hemiptera: Pseudococcidae). Ecology and evolution, 9(4):1972–1984.

Liu, Y., Shen, Z., Yu, J., Li, Z., Liu, X., and Xu, H. (2020). Comparison of gut bacterial communities and their associations with host diets in four fruit borers. Pest management science, 76(4):1353–1362.

Losey, J. E. and Vaughan, M. (2006). The economic value of ecological services provided by insects. Bioscience, 56(4):311–323.

Lutz, H. L., Jackson, E. W., Webala, P. W., Babyesiza, W. S., Peterhans, J. C. K., Demos, T. C., Patterson, B. D., and Gilbert, J. A. (2019). Ecology and host identity outweigh evolutionary history in shaping the bat microbiome. Msystems, 4(6).

Magoč, T. and Salzberg, S. L. (2011). Flash: fast length adjustment of short reads to improve genome assemblies. Bioinformatics, 27(21):2957–2963.

Maillet, A., Bouju-Albert, A., Roblin, S., Vaissié, P., Leuillet, S., Dousset, X., Jaffrès, E., Combrisson, J., and Prévost, H. (2021). Impact of dna extraction and sampling methods on bacterial communities monitored by 16s rdna metabarcoding in cold-smoked salmon and processing plant surfaces. Food Microbiology, 95:103705.

Majumder, R., Sutcliffe, B., Taylor, P. W., and Chapman, T. A. (2019). Next-generation sequencing reveals relationship between the larval microbiome and food substrate in the polyphagous queensland fruit fly. Scientific reports, 9(1):1–12.

Malacrinò, A. (2018). Meta-omics tools in the world of insect-microorganism interactions. Biology, 7:50.

Malacrinò, A., Campolo, O., Medina, R. F., and Palmeri, V. (2018). Instar-and host-associated differentiation of bacterial communities in the mediterranean fruit fly ceratitis capitata. PloS one, 13(3):e0194131.

Mancini, M., Damiani, C., Accoti, A., Tallarita, M., Nunzi, E., Cappelli, A., Bozic, J., Catanzani, R., Rossi, P., Valzano, M., et al. (2018). Estimating bacteria diversity in different organs of nine species of mosquito by next generation sequencing. BMC microbiology, 18(1):1–10.

Martinez, A. J., Onchuru, T. O., Ingham, C. S., Sandoval-Calderón, M., Salem, H., Deckert, J., and Kaltenpoth, M. (2019). Angiosperm to gymnosperm host-plant switch entails shifts in microbiota of the welwitschia bug, probergrothius angolensis (distant, 1902). Molecular ecology, 28(23):5172–5187.

Mason, C. J., Campbell, A. M., Scully, E. D., and Hoover, K. (2019). Bacterial and fungal midgut community dynamics and transfer between mother and brood in the asian longhorned beetle (anoplophora glabripennis), an invasive xylophage. Microbial ecology, 77(1):230–242.

Mazel, F., Davis, K. M., Loudon, A., Kwong, W. K., Groussin, M., and Parfrey, L. W. (2018). Is host filtering the main driver of phylosymbiosis across the tree of life? Msystems, 3(5).

McLean, A. H., Godfray, H. C. J., Ellers, J., and Henry, L. M. (2019). Host relatedness influences the composition of aphid microbiomes. Environmental microbiology reports, 11(6):808–816.

McMurdie, P. J. and Holmes, S. (2013). phyloseq: an r package for reproducible interactive analysis and graphics of microbiome census data. PloS one, 8(4):e61217.

McOrist, A. L., Jackson, M., and Bird, A. R. (2002). A comparison of five methods for extraction of bacterial dna from human faecal samples. Journal of microbiological methods, 50(2):131–139.

Meng, F., Bar-Shmuel, N., Shavit, R., Behar, A., and Segoli, M. (2019a). Gut bacteria of weevils developing on plant roots under extreme desert conditions. BMC microbiology, 19(1):1–13.

Meng, L., Li, X., Cheng, X., and Zhang, H. (2019b). 16s rrna gene sequencing reveals a shift in the microbiota of diaphorina citri during the psyllid life cycle. Frontiers in microbiology, 10:1948.

Mereghetti, V., Chouaia, B., Limonta, L., Locatelli, D. P., and Montagna, M. (2019). Evidence for a conserved microbiota across the different developmental stages of plodia interpunctella. Insect science, 26(3):466–478.

Mikaelyan, A., Dietrich, C., Köhler, T., Poulsen, M., Sillam-Dussès, D., and Brune, A. (2015). Diet is the primary determinant of bacterial community structure in the guts of higher termites. Molecular ecology, 24(20):5284–5295.

Minard, G., Tikhonov, G., Ovaskainen, O., and Saastamoinen, M. (2019). The microbiome of the melitaea cinxia butterfly shows marked variation but is only little explained by the traits of the butterfly or its host plant. Environmental microbiology, 21(11):4253–4269.

Montagna, M., Mereghetti, V., Gargari, G., Guglielmetti, S., Faoro, F., Lozzia, G., Locatelli, D., and Limonta, L. (2016). Evidence of a bacterial core in the stored products pest plodia interpunctella: the influence of different diets. Environmental Microbiology, 18(12):4961–4973.

Nobles, S. and Jackson, C. R. (2020). Effects of life stage, site, and species on the dragonfly gut microbiome. Microorganisms, 8(2):183.

Parada, A. E., Needham, D. M., and Fuhrman, J. A. (2016). Every base matters: assessing small subunit rrna primers for marine microbiomes with mock communities, time series and global field samples. Environmental Microbiology, 18(5):1403–1414.

Paradis, E. and Schliep, K. (2019). ape 5.0: an environment for modern phylogenetics and evolutionary analyses in r. Bioinformatics, 35(3):526–528.

Parfrey, L. W., Moreau, C. S., and Russell, J. A. (2018). Introduction: The host-associated microbiome: Pattern, process and function. Molecular ecology, 27(8):1749–1765.

Parr, C. S. and Cummings, M. P. (2005). Data sharing in ecology and evolution. Trends in Ecology & Evolution, 20(7):362–363.

Pennington, M. J., Prager, S. M., Walton, W. E., and Trumble, J. T. (2016). Culex quinquefasciatus larval microbiomes vary with instar and exposure to common wastewater contaminants. Scientific reports, 6(1):1–9.

Pérez-Cobas, A. E., Maiques, E., Angelova, A., Carrasco, P., Moya, A., and Latorre, A. (2015). Diet shapes the gut microbiota of the omnivorous cockroach blattella germanica. FEMS microbiology ecology, 91(4):fiv022.

Phalnikar, K., Kunte, K., and Agashe, D. (2018). Dietary and developmental shifts in butterfly-associated bacterial communities. Royal Society open science, 5(5):171559.

Quast, C., Pruesse, E., Yilmaz, P., Gerken, J., Schweer, T., Yarza, P., Peplies, J., and Glöckner, F. O. (2012). The silva ribosomal rna gene database project: improved data processing and web-based tools. Nucleic acids research, 41(D1):D590–D596.

R Core Team (2020). R: A language and environment for statistical computing. version 4.0. 2 (taking off again). R Foundation for Statistical Computing, Vienna, Austria.

Ramalho, M. O., Bueno, O. C., and Moreau, C. S. (2017). Microbial composition of spiny ants (hymenoptera: Formicidae: Polyrhachis) across their geographic range. BMC Evolutionary Biology, 17(1):1–15.

Rassati, D., Marini, L., and Malacrinò, A. (2019). Acquisition of fungi from the environment modifies ambrosia beetle mycobiome during invasion. PeerJ, 7:e8103.

Raymann, K., Motta, E. V., Girard, C., Riddington, I. M., Dinser, J. A., and Moran, N. A. (2018). Imidacloprid decreases honey bee survival rates but does not affect the gut microbiome. Applied and environmental microbiology, 84(13).

Receveur, J. P., Pechal, J. L., Benbow, M. E., Donato, G., Rainey, T., and Wallace, J. R. (2018). Changes in larval mosquito microbiota reveal non-target effects of insecticide treatments in hurricane-created habitats. Microbial ecology, 76(3):719–728.

Roche, D. G., Kruuk, L. E., Lanfear, R., and Binning, S. A. (2015). Public data archiving in ecology and evolution: how well are we doing? PLoS Biol, 13(11):e1002295.

Rodríguez-Ruano, S. M., Škochová, V., Rego, R. O., Schmidt, J. O., Roachell, W., Hypša, V., and Nováková, E. (2018). Microbiomes of north american triatominae: the grounds for chagas disease epidemiology. Frontiers in microbiology, 9:1167.

Rognes, T., Flouri, T., Nichols, B., Quince, C., and Mahé, F. (2016). Vsearch: a versatile open source tool for metagenomics. PeerJ, 4:e2584.

Roth-Schulze, A. J., Pintado, J., Zozaya-Valdés, E., Cremades, J., Ruiz, P., Kjelleberg, S., and Thomas, T. (2018). Functional biogeography and host specificity of bacterial communities associated with the marine green alga ulva spp. Molecular ecology, 27(8):1952–1965.

Rothman, J. A., Leger, L., Graystock, P., Russell, K., and McFrederick, Q. S. (2019a). The bumble bee microbiome increases survival of bees exposed to selenate toxicity. Environmental microbiology, 21(9):3417–3429.

Rothman, J. A., Leger, L., Kirkwood, J. S., and McFrederick, Q. S. (2019b). Cadmium and selenate exposure affects the honey bee microbiome and metabolome, and bee-associated bacteria show potential for bioaccumulation. Applied and environmental microbiology, 85(21).

Russell, J. A., Moreau, C. S., Goldman-Huertas, B., Fujiwara, M., Lohman, D. J., and Pierce, N. E. (2009). Bacterial gut symbionts are tightly linked with the evolution of herbivory in ants. Proceedings of the National Academy of Sciences, 106(50):21236–21241.

Segata, N., Baldini, F., Pompon, J., Garrett, W. S., Truong, D. T., Dabiré, R. K., Diabaté, A., Levashina, E. A., and Catteruccia, F. (2016). The reproductive tracts of two malaria vectors are populated by a core microbiome and by gender-and swarm-enriched microbial biomarkers. Scientific reports, 6(1):1–10.

Skrodenytė-Arbačiauskienė, V., Budrienė, A., Blažytė-Čereškienė, L., and Budrys, E. (2019). Illumina-based 16s metagenomic analysis of the indigenous gut microbiota of cavity-nesting bee megachile centuncularis: a comparison with the cavity-nesting wasp ancistrocerus antilope. Journal of Apicultural Research, 58(4):587–590.

Stathopoulou, P., Asimakis, E. D., Khan, M., Caceres, C., Bourtzis, K., and Tsiamis, G. (2019). Irradiation effect on the structure of bacterial communities associated with the oriental fruit fly, bactrocera dorsalis. Entomologia Experimentalis et Applicata, 167(3):209–219.

Staudacher, H., Kaltenpoth, M., Breeuwer, J. A., Menken, S. B., Heckel, D. G., and Groot, A. T. (2016). Variability of bacterial communities in the moth heliothis virescens indicates transient association with the host. PLoS One, 11(5):e0154514.

Strano, C. P., Malacrinò, A., Campolo, O., and Palmeri, V. (2018). Influence of host plant on thaumetopoea pityocampa gut bacterial community. Microbial ecology, 75(2):487–494.

Suenami, S., Nobu, M. K., and Miyazaki, R. (2019). Community analysis of gut microbiota in hornets, the largest eusocial wasps, vespa mandarinia and v. simillima. Scientific reports, 9(1):1–13.

Sun, T., Wang, X.-Q., Zhao, Z.-L., Yu, S.-H., Yang, P., and Chen, X.-M. (2018). A lethal fungus infects the chinese white wax scale insect and causes dramatic changes in the host microbiota. Scientific reports, 8(1):1–8.

Tinker, K. A. and Ottesen, E. A. (2016). The core gut microbiome of the american cockroach, periplaneta americana, is stable and resilient to dietary shifts. Applied and Environmental Microbiology, 82(22):6603–6610.

Tremblay, J., Singh, K., Fern, A., Kirton, E., He, S., Woyke, T., Lee, J., Chen, F., Dangl, J., and Tringe, S. (2015). Primer and platform effects on 16s rrna tag sequencing. Frontiers in Microbiology, 6:771.

Trevelline, B. K., Fontaine, S. S., Hartup, B. K., and Kohl, K. D. (2019). Conservation biology needs a microbial renaissance: a call for the consideration of host-associated microbiota in wildlife management practices. Proceedings of the Royal Society B, 286(1895):20182448.

Trevelline, B. K., Sosa, J., Hartup, B. K., and Kohl, K. D. (2020). A bird’s-eye view of phylosymbiosis: weak signatures of phylosymbiosis among all 15 species of cranes. Proceedings of the Royal Society B, 287(1923):20192988.

Vacheron, J., Péchy-Tarr, M., Brochet, S., Heiman, C. M., Stojiljkovic, M., Maurhofer, M., and Keel, C. (2019). T6ss contributes to gut microbiome invasion and killing of an herbivorous pest insect by plant-beneficial pseudomonas protegens. The ISME journal, 13(5):1318–1329.

van Schooten, B., Godoy-Vitorino, F., McMillan, W. O., and Papa, R. (2018). Conserved microbiota among young heliconius butterfly species. PeerJ, 6:e5502.

Vasselon, V., Domaizon, I., Rimet, F., Kahlert, M., and Bouchez, A. (2017). Application of high-throughput sequencing (hts) metabarcoding to diatom biomonitoring: Do dna extraction methods matter? Freshwater Science, 36(1):162–177.

Ventura, C., Briones-Roblero, C. I., Hernández, E., Rivera-Orduña, F. N., and Zúñiga, G. (2018). Comparative analysis of the gut bacterial community of four anastrepha fruit flies (diptera: Tephritidae) based on pyrosequencing. Current microbiology, 75(8):966–976.

Walker, A. W., Martin, J. C., Scott, P., Parkhill, J., Flint, H. J., and Scott, K. P. (2015). 16s rrna gene-based profiling of the human infant gut microbiota is strongly influenced by sample processing and pcr primer choice. Microbiome, 3(1):1–11.

Waltmann, A., Willcox, A. C., Balasubramanian, S., Mayori, K. B., Guerrero, S. M., Sanchez, R. S. S., Roach, J., Pino, C. C., Gilman, R. H., Bern, C., et al. (2019). Hindgut microbiota in laboratory-reared and wild triatoma infestans. PLoS neglected tropical diseases, 13(5):e0007383.

Wang, X., Gao, Q., Wang, W., Wang, X., Lei, C., and Zhu, F. (2018). The gut bacteria across life stages in the synanthropic fly chrysomya megacephala. BMC microbiology, 18(1):1–8.

Whitaker, M. R., Salzman, S., Sanders, J., Kaltenpoth, M., and Pierce, N. E. (2016). Microbial communities of lycaenid butterflies do not correlate with larval diet. Frontiers in Microbiology, 7:1920.

Whitlock, M. C. (2011). Data archiving in ecology and evolution: best practices. Trends in ecology & evolution, 26(2):61–65.

Wilches, D., Laird, R., Fields, P., Coghlin, P., and Floate, K. (2018). Spiroplasma dominates the microbiome of khapra beetle: comparison with a congener, effects of life stage and temperature. Symbiosis, 76(3):277–291.

Wohlfahrt, D., Woolf, M. S., and Singh, B. (2020). A survey of bacteria associated with various life stages of primary colonizers: Lucilia sericata and phormia regina. Science & Justice, 60(2):173–179.

Woruba, D. N., Morrow, J. L., Reynolds, O. L., Chapman, T. A., Collins, D. P., and Riegler, M. (2019). Diet and irradiation effects on the bacterial community composition and structure in the gut of domesticated teneral and mature queensland fruit fly, bactrocera tryoni (diptera: Tephritidae). BMC microbiology, 19(1):1–13.

Wu, H.-X., Chen, X., Chen, H., Lu, Q., Yang, Z., Ren, W., Liu, J., Shao, S., Wang, C., King-Jones, K., et al. (2018). Variation and diversification of the microbiome of schlechtendalia chinensis on two alternate host plants. PloS one, 13(11):e0200049.

Xu, L., Lu, M., Xu, D., Chen, L., and Sun, J. (2016). Sexual variation of bacterial microbiota of dendroctonus valens guts and frass in relation to verbenone production. Journal of insect physiology, 95:110–117.

Xu, S., Jiang, L., Qiao, G., and Chen, J. (2019). The bacterial flora associated with the polyphagous aphid aphis gossypii glover (hemiptera: Aphididae) is strongly affected by host plants. Microbial ecology, pages 1–14.

Xu, T.-T., Jiang, L.-Y., Chen, J., and Qiao, G.-X. (2020). Host plants influence the symbiont diversity of eriosomatinae (hemiptera: Aphididae). Insects, 11(4):217.

Xue, Z., Zhang, J., Zhang, R., Huang, Z., Wan, Q., and Zhang, Z. (2019). Comparative analysis of gut bacterial communities in housefly larvae fed different diets using a high-throughput sequencing approach. FEMS microbiology letters, 366(11):fnz126.

Yang, L. H. and Gratton, C. (2014). Insects as drivers of ecosystem processes. Current Opinion in Insect Science, 2:26–32.

Yao, Z., Ma, Q., Cai, Z., Raza, M. F., Bai, S., Wang, Y., Zhang, P., Ma, H., and Zhang, H. (2019). Similar shift patterns in gut bacterial and fungal communities across the life stages of bactrocera minax larvae from two field populations. Frontiers in microbiology, 10:2262.

Yong, H.-S., Song, S.-L., Chua, K.-O., and Lim, P.-E. (2017). Microbiota associated with bactrocera carambolae and b. dorsalis (insecta: Tephritidae) revealed by next-generation sequencing of 16s rrna gene. Meta Gene, 11:189–196.

Yun, J.-H., Roh, S. W., Whon, T. W., Jung, M.-J., Kim, M.-S., Park, D.-S., Yoon, C., Nam, Y.-D., Kim, Y.-J., Choi, J.-H., et al. (2014). Insect gut bacterial diversity determined by environmental habitat, diet, developmental stage, and phylogeny of host. Applied and Environmental Microbiology, 80(17):5254–5264.

Zepeda-Paulo, F., Ortiz-Martínez, S., Silva, A. X., and Lavandero, B. (2018). Low bacterial community diversity in two introduced aphid pests revealed with 16s rrna amplicon sequencing. PeerJ, 6:e4725.

Zhang, F., Sun, X. X., Zhang, X. C., Zhang, S., Lu, J., Xia, Y. M., Huang, Y. H., and Wang, X. J. (2018a). The interactions between gut microbiota and entomopathogenic fungi: a potential approach for biological control of blattella germanica (l.). Pest management science, 74(2):438–447.

Zhang, F. and Yang, R. (2019). Life history and functional capacity of the microbiome are altered in beta-cypermethrin-resistant cockroaches. International journal for parasitology, 49(9):715–723.

Zhang, Z., Jiao, S., Li, X., and Li, M. (2018b). Bacterial and fungal gut communities of agrilus mali at different developmental stages and fed different diets. Scientific reports, 8(1):1–11.

Zhao, C., Zhao, H., Zhang, S., Luo, J., Zhu, X., Wang, L., Zhao, P., Hua, H., and Cui, J. (2019). The developmental stage symbionts of the pea aphid-feeding chrysoperla sinica (tjeder). Frontiers in microbiology, 10:2454.

Zheng, Y., Xiao, G., Zhou, W., Gao, Y., Li, Z., Du, G., and Chen, B. (2020). Midgut microbiota diversity of potato tuber moth associated with potato tissue consumed. BMC microbiology, 20(1):1–16.

Zhu, D., Chen, Q.-L., Li, H., Yang, X.-R., Christie, P., Ke, X., and Zhu, Y.-G. (2018a). Land use influences antibiotic resistance in the microbiome of soil collembolans orchesellides sinensis. Environmental science & technology, 52(24):14088–14098.

Zhu, D., Zheng, F., Chen, Q.-L., Yang, X.-R., Christie, P., Ke, X., and Zhu, Y.-G. (2018b). Exposure of a soil collembolan to ag nanoparticles and agno3 disturbs its associated microbiota and lowers the incidence of antibiotic resistance genes in the gut. Environmental science & technology, 52(21):12748–12756.

