## Supplementary Material 2 for "Host species identity shapes the diversity and structure of insect microbiota": SM2.html


Code 

- Show All Code
- Hide All Code
- Download Rmd

### Supplementary Material 2


#### **Tab. 2.1** — DNA extraction method

#### **Tab. 2.2** — Primers

#### **Fig. 2.1** — Primers

#### **Tab. 2.3** — 16S Region

#### **Tab. 2.4** — Sequencing technology

#### **Tab. 2.5** — Sequencing platform

#### **Tab. 2.6** — Database

#### **Tab. 2.7** — Insect ID

#### **Tab. 2.8** — Search

#### **Tab. 2.9** — Comment
