## Supplementary Material 3 for "Host species identity shapes the diversity and structure of insect microbiota": SM3.html


Code 

- Show All Code
- Hide All Code
- Download Rmd

### Supplementary Material 3


#### **Tab. 3.1** — Insect species

#### **Tab. 3.2** — Insect order

#### **Tab. 3.3** — Sex

#### **Tab. 3.4** — Life stage

#### **Tab. 3.5** — Sample origin

#### **Tab. 3.6** — Diet

#### **Tab. 3.7** — Sample tissue

#### **Tab. 3.8** — Sample treatment
