## Supplementary Material 4 for "Host species identity shapes the diversity and structure of insect microbiota": SM4.pdf

### Methods

Paired-end reads were merged using FLASH 1.2.11 [1] and data were processed using the DADA2 [2] pipeline implemented in R to remove low-quality data, identify ASVs and remove chimera. Taxonomy was assigned using SILVA v138 [3]. Reads/samples filtering and downstream analyses performed as indicated in the main text.

### Results

Due to the more stringent filtering of the DADA2 pipeline, the analyses were performed on 968 samples for whole insects (including 66 species from the orders Coleoptera, Diptera, Lepidoptera, and Hymenoptera) and 1,018 for insect guts (including 54 species from the orders Hymenoptera, Lepidoptera, Blattodea, and Diptera).

Similarly to the results obtained using the VSEARCH pipeline, insect species explained most of the variation in both whole insects (0.39-0.81%) and insect guts (0.34-0.86%). while the other factors explained $\leq$ 1.5% of the variance (Tab. S4.1). Conversely, it is possible to notice the microbial diversity was significantly influenced by insect species and origin, but not from the other factors. Similarly, the factor treatment did not had a significant effect on the microbiota of whole insects and insect guts.

**Tab. S4.1**. Analysis of the effect of insect species, sex, life stage, diet, sample treatment, and sample origin on the structure and diversity of insect-associated bacterial communities. The effect of each factor on the structure of microbial communities was tested using a PERMANOVA on a Bray-Curtis (accounting for relative abundance) or Jaccard (accounting for presence/absence) distance matrix. The effect of factors on microbial diversity was tested using a linear mixed-effects model on Shannon diversity index. Results are grouped by whole insect specimens (upper) or insect guts (lower).

| | | **Bray-Curtis** | | | **Jaccard** | | | **Shannon** | | |
|---|---|---|---|---|---|---|---|---|---|---|
| **Factors (whole insect)** | df | $R^2$ | F | $P$ | $R^2$ | F | $P$ | $R^2$ | $\chi^2$ | $P$ |
| Insect species | 65 | 0.49 | 14.57 | <0.001 | 0.39 | 9.35 | <0.001 | 0.81 | 285.1 | <0.001 |
| Sex | 2 | 0.004 | 4.74 | <0.001 | 0.004 | 3.74 | <0.001 | 0.002 | 1.47 | 0.4 |
| Life stage | 3 | 0.01 | 9.07 | <0.001 | 0.01 | 6.5 | <0.001 | 0.001 | 2.99 | 0.4 |
| Diet | 2 | 0.006 | 6.28 | <0.001 | 0.006 | 4.93 | <0.001 | 0.05 | 0.3 | 0.8 |
| Treatment | 1 | 0.0006 | 1.23 | 0.058 | 0.0008 | 1.23 | 0.03 | 0.0009 | 2.56 | 0.1 |
| Origin | 1 | 0.006 | 12.21 | <0.001 | 0.006 | 9.67 | <0.001 | 0.06 | 17.35 | <0.001 |
| **Factors (guts)** | df | $R^2$ | F | $P$ | $R^2$ | F | $P$ | $R^2$ | $\chi^2$ | $P$ |
| Insect species | 53 | 0.46 | 17.17 | <0.001 | 0.34 | 10.44 | <0.001 | 0.86 | 2474.5 | <0.001 |
| Sex | 2 | 0.004 | 4.17 | <0.001 | 0.003 | 3.06 | <0.001 | 0.004 | 9.89 | 0.007 |
| Life stage | 2 | 0.001 | 1.08 | 0.56 | 0.001 | 1.04 | 0.54 | 0.001 | 1.73 | 0.4 |
| Diet | 2 | 0.01 | 10.35 | <0.001 | 0.008 | 6.78 | <0.001 | 0.01 | 4.15 | 0.1 |
| Treatment | 1 | 0.001 | 2.26 | 0.058 | 0.001 | 2.29 | 0.02 | 0.0001 | 0.4 | 0.5 |
| Origin | 1 | 0.03 | 76.26 | <0.001 | 0.03 | 49.58 | <0.001 | 0.03 | 24.72 | <0.001 |

When testing for a phylogenetic signal in the microbial community diversity (Shannon index) and structure (Bray-Curtis distance) on insect guts ($n$=32), results ($\lambda$ = 0.938) suggest that bacterial diversity in insect guts is non-random ($\lambda \neq 0$, $P < 0.001$) and shows weak phylogenetic signal ($\lambda \neq 1$,

$P < 0.001$). These results were confirmed by the Blomberg's $K$ test ($K = 0.16$, $P = 0.17$). Similarly, the Mantel's test does not show a significant correlation between the structure of microbiota and the host phylogenetic distance ($r = 0.04$, $P = 0.17$).

In addition, there are many similarities in the differences in relative abundance between taxa among different groups. For example, *Enterobacter* has been found to be more abundant in juveniles than adults, pupae and eggs of whole insects ($\chi^2$=79.59, $P$<0.001). Also, *Pseudomonas* was more abundant in insects reared in the lab than those collected in the field ($\chi^2$=34.57, $P$<0.001). *Wolbachia* was more abundant in carnivores than herbivores or insects feeding on artificial diet. In insect guts, *Alistipes*, *Bacteroides*, *Desulfovibrio*, *Christensenellaceae R-7 group* were consistently more abundant ($P < 0.05$) in Blattodea than Diptera, Hymenoptera and Lepidoptera. These are just few example showing the overlap between the two approaches used in this study. Unfortunately, the differences in the quality-control procedures and data processing remove a different numbers of samples, so we cannot directly compare the two approaches. However, this overlap in the taxa that show significant change in the same direction when comparing different groups suggest that results hold regardless of the approach (VSEARCH vs DADA2, OTUs vs ASVs) or version of the database used for taxonomical annotation (SILVA v132 vs v138).

### References

[1] Magoč, T., & Salzberg, S. L. (2011). FLASH: fast length adjustment of short reads to improve genome assemblies. Bioinformatics, 27(21), 2957-2963.

[2] Callahan, B. J., McMurdie, P. J., Rosen, M. J., Han, A. W., Johnson, A. J. A., & Holmes, S. P. (2016). DADA2: high-resolution sample inference from Illumina amplicon data. Nature methods, 13(7), 581-583.

[3] Quast, C., Pruesse, E., Yilmaz, P., Gerken, J., Schweer, T., Yarza, P., ... & Glöckner, F. O. (2012). The SILVA ribosomal RNA gene database project: improved data processing and web-based tools. Nucleic acids research, 41(D1), D590-D596.
