## Supplementary figures and images for "Host species identity shapes the diversity and structure of insect microbiota"

### SM5.pdf

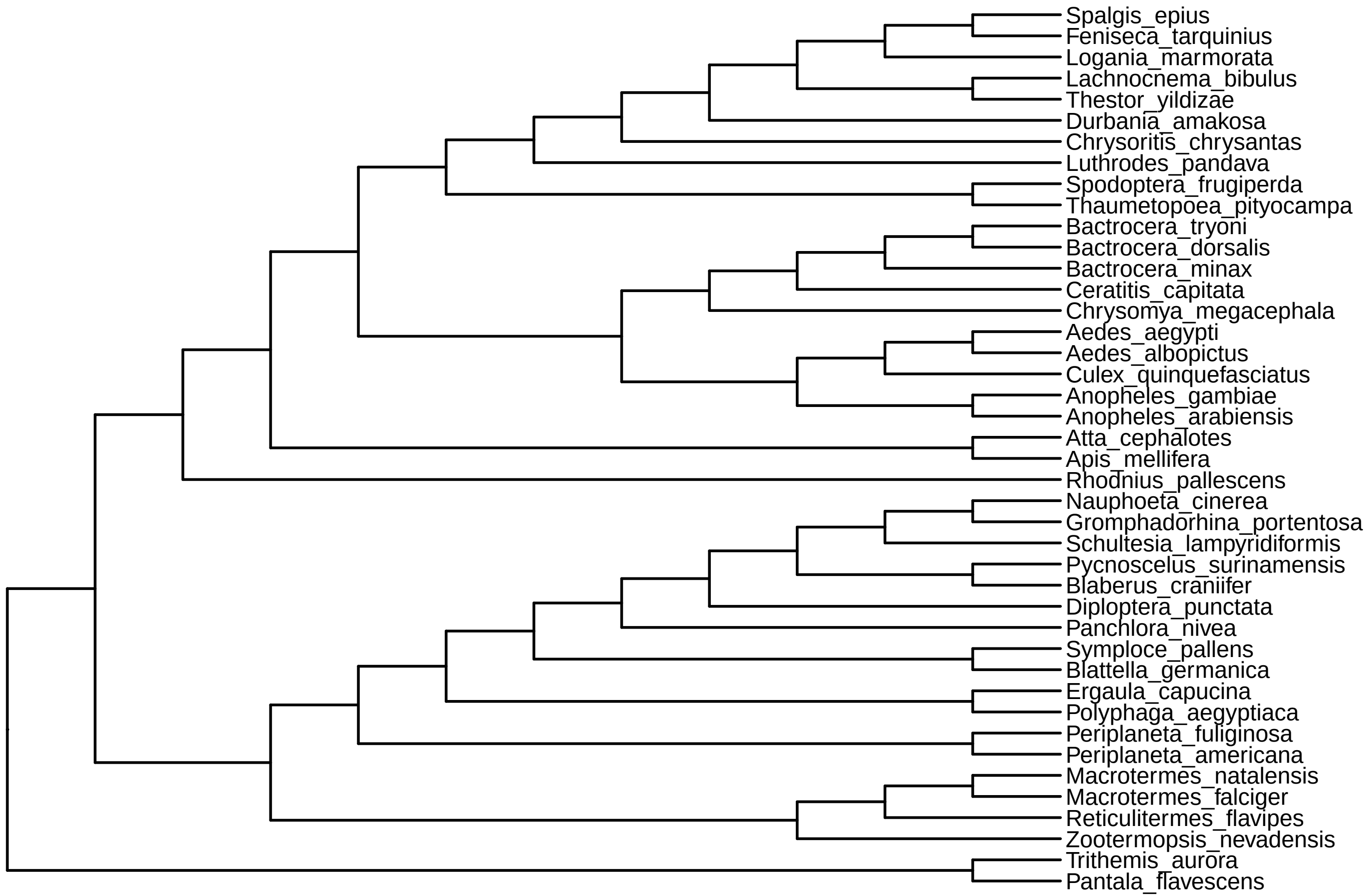
