## Supplementary Material 6 for "Host species identity shapes the diversity and structure of insect microbiota": SM6.html


Code 

- Show All Code
- Hide All Code
- Download Rmd

### Supplementary material 6

### Differentially abundant taxa - Whole


#### Insect order

##### Model

##### Posthoc


```
#### Acinetobacter 
 contrast                  estimate     SE   df t.ratio p.value
 coleoptera - diptera       0.03223 0.0140 26.1   2.307  0.1746
 coleoptera - hemiptera     0.04452 0.0154 28.9   2.899  0.0510
 coleoptera - hymenoptera   0.02818 0.0166 26.3   1.695  0.4542
 coleoptera - lepidoptera   0.03960 0.0160 24.7   2.478  0.1285
 diptera - hemiptera        0.01228 0.0151 29.6   0.815  0.9239
 diptera - hymenoptera     -0.00405 0.0166 24.1  -0.245  0.9991
 diptera - lepidoptera      0.00737 0.0157 25.1   0.469  0.9895
 hemiptera - hymenoptera   -0.01633 0.0177 26.2  -0.921  0.8862
 hemiptera - lepidoptera   -0.00491 0.0170 27.4  -0.290  0.9984
 hymenoptera - lepidoptera  0.01142 0.0183 23.4   0.625  0.9696

Degrees-of-freedom method: kenward-roger 
P value adjustment: tukey method for comparing a family of 5 estimates 

 

#### Lactobacillus 
 contrast                   estimate     SE   df t.ratio p.value
 coleoptera - diptera       0.004333 0.0212 28.8   0.204  0.9996
 coleoptera - hemiptera     0.005714 0.0230 31.2   0.248  0.9991
 coleoptera - hymenoptera  -0.092674 0.0250 35.7  -3.709  0.0060
 coleoptera - lepidoptera   0.005849 0.0245 26.2   0.239  0.9992
 diptera - hemiptera        0.001381 0.0225 31.4   0.061  1.0000
 diptera - hymenoptera     -0.097007 0.0252 29.5  -3.846  0.0050
 diptera - lepidoptera      0.001516 0.0241 26.2   0.063  1.0000
 hemiptera - hymenoptera   -0.098388 0.0268 31.2  -3.673  0.0074
 hemiptera - lepidoptera    0.000135 0.0257 28.1   0.005  1.0000
 hymenoptera - lepidoptera  0.098522 0.0281 27.3   3.509  0.0126

Degrees-of-freedom method: kenward-roger 
P value adjustment: tukey method for comparing a family of 5 estimates 

 

#### Saccharibacter 
 contrast                  estimate     SE   df t.ratio p.value
 coleoptera - diptera       0.00279 0.0139 29.5   0.200  0.9996
 coleoptera - hemiptera     0.00279 0.0151 31.6   0.185  0.9997
 coleoptera - hymenoptera  -0.04440 0.0163 39.1  -2.730  0.0674
 coleoptera - lepidoptera   0.00279 0.0161 26.7   0.173  0.9998
 diptera - hemiptera        0.00000 0.0147 31.6   0.000  1.0000
 diptera - hymenoptera     -0.04719 0.0165 31.1  -2.856  0.0545
 diptera - lepidoptera      0.00000 0.0158 26.5   0.000  1.0000
 hemiptera - hymenoptera   -0.04719 0.0175 32.6  -2.694  0.0767
 hemiptera - lepidoptera    0.00000 0.0169 28.3   0.000  1.0000
 hymenoptera - lepidoptera  0.04719 0.0184 28.4   2.558  0.1058

Degrees-of-freedom method: kenward-roger 
P value adjustment: tukey method for comparing a family of 5 estimates
```

#### Sex

##### Model

##### Posthoc


```
#### Arsenophonus 
 contrast         estimate     SE   df t.ratio p.value
 female - male     -0.0493 0.0175 2426  -2.813  0.0137
 female - unknown  -0.0418 0.0108 2371  -3.876  0.0003
 male - unknown     0.0075 0.0186 2439   0.404  0.9139

Degrees-of-freedom method: kenward-roger 
P value adjustment: tukey method for comparing a family of 3 estimates 

 

#### Salinibacter 
 contrast         estimate     SE   df t.ratio p.value
 female - male      0.0115 0.0177 2429   0.651  0.7918
 female - unknown   0.0299 0.0105 1200   2.856  0.0121
 male - unknown     0.0183 0.0186 2237   0.987  0.5851

Degrees-of-freedom method: kenward-roger 
P value adjustment: tukey method for comparing a family of 3 estimates 

 

#### Wolbachia 
 contrast         estimate     SE   df t.ratio p.value
 female - male     -0.0518 0.0293 2421  -1.768  0.1805
 female - unknown   0.0494 0.0181 2419   2.733  0.0174
 male - unknown     0.1012 0.0311 2435   3.258  0.0033

Degrees-of-freedom method: kenward-roger 
P value adjustment: tukey method for comparing a family of 3 estimates
```

#### Life stage

##### Model

##### Posthoc


```
#### Arsenophonus 
 contrast           estimate      SE     df t.ratio p.value
 adult - egg        -0.00887 0.02465 2434.5  -0.360  0.9964
 adult - juvenile   -0.02923 0.00933 2387.2  -3.132  0.0151
 adult - pupae      -0.01661 0.01945 2420.5  -0.854  0.9134
 adult - unknown     0.01253 0.09451   34.3   0.133  0.9999
 egg - juvenile     -0.02037 0.02526 2436.0  -0.806  0.9288
 egg - pupae        -0.00774 0.03032 2428.8  -0.255  0.9991
 egg - unknown       0.02139 0.09742   38.7   0.220  0.9995
 juvenile - pupae    0.01262 0.01909 2415.9   0.661  0.9645
 juvenile - unknown  0.04176 0.09461   34.5   0.441  0.9918
 pupae - unknown     0.02914 0.09622   36.9   0.303  0.9981

Degrees-of-freedom method: kenward-roger 
P value adjustment: tukey method for comparing a family of 5 estimates 

 

#### Wolbachia 
 contrast           estimate     SE     df t.ratio p.value
 adult - egg         0.11575 0.0412 2429.2   2.811  0.0399
 adult - juvenile    0.03917 0.0156 2424.7   2.506  0.0896
 adult - pupae       0.00851 0.0325 2416.5   0.262  0.9990
 adult - unknown     0.12831 0.1859   34.0   0.690  0.9572
 egg - juvenile     -0.07658 0.0422 2431.5  -1.815  0.3652
 egg - pupae        -0.10724 0.0506 2423.4  -2.118  0.2125
 egg - unknown       0.01256 0.1900   37.1   0.066  1.0000
 juvenile - pupae   -0.03066 0.0319 2412.8  -0.962  0.8719
 juvenile - unknown  0.08914 0.1860   34.1   0.479  0.9888
 pupae - unknown     0.11980 0.1883   35.8   0.636  0.9681

Degrees-of-freedom method: kenward-roger 
P value adjustment: tukey method for comparing a family of 5 estimates 

 

#### Enterococcus 
 contrast            estimate      SE     df t.ratio p.value
 adult - egg         0.000931 0.01391 2412.7   0.067  1.0000
 adult - juvenile   -0.003984 0.00531 2427.0  -0.751  0.9444
 adult - pupae      -0.049432 0.01095 2407.8  -4.512  0.0001
 adult - unknown     0.041355 0.11817   33.3   0.350  0.9966
 egg - juvenile     -0.004915 0.01427 2413.9  -0.345  0.9970
 egg - pupae        -0.050363 0.01710 2410.2  -2.946  0.0269
 egg - unknown       0.040424 0.11892   34.1   0.340  0.9970
 juvenile - pupae   -0.045448 0.01075 2406.6  -4.228  0.0002
 juvenile - unknown  0.045339 0.11820   33.3   0.384  0.9952
 pupae - unknown     0.090787 0.11861   33.8   0.765  0.9387

Degrees-of-freedom method: kenward-roger 
P value adjustment: tukey method for comparing a family of 5 estimates 

 

#### Leucobacter 
 contrast            estimate      SE     df t.ratio p.value
 adult - egg        -3.38e-02 0.01782 2421.4  -1.898  0.3185
 adult - juvenile   -1.05e-01 0.00667 1992.6 -15.737  <.0001
 adult - pupae      -6.04e-02 0.01411 2433.1  -4.283  0.0002
 adult - unknown    -3.38e-02 0.04570   35.9  -0.740  0.9456
 egg - juvenile     -7.11e-02 0.01825 2409.2  -3.898  0.0009
 egg - pupae        -2.66e-02 0.02196 2436.1  -1.211  0.7450
 egg - unknown       2.67e-05 0.04879   46.4   0.001  1.0000
 juvenile - pupae    4.45e-02 0.01386 2427.0   3.214  0.0116
 juvenile - unknown  7.12e-02 0.04581   36.3   1.553  0.5358
 pupae - unknown     2.66e-02 0.04755   42.1   0.560  0.9801

Degrees-of-freedom method: kenward-roger 
P value adjustment: tukey method for comparing a family of 5 estimates 

 

#### Escherichia-Shigella 
 contrast            estimate      SE     df t.ratio p.value
 adult - egg        -0.012064 0.01486 2427.0  -0.812  0.9270
 adult - juvenile   -0.022290 0.00565 2431.4  -3.948  0.0008
 adult - pupae      -0.012841 0.01171 2415.1  -1.097  0.8084
 adult - unknown     0.003788 0.07160   33.9   0.053  1.0000
 egg - juvenile     -0.010226 0.01523 2429.3  -0.672  0.9625
 egg - pupae        -0.000777 0.01827 2421.4  -0.043  1.0000
 egg - unknown       0.015852 0.07300   36.6   0.217  0.9995
 juvenile - pupae    0.009450 0.01149 2411.8   0.822  0.9238
 juvenile - unknown  0.026079 0.07165   34.0   0.364  0.9961
 pupae - unknown     0.016629 0.07242   35.4   0.230  0.9994

Degrees-of-freedom method: kenward-roger 
P value adjustment: tukey method for comparing a family of 5 estimates 

 

#### Leuconostoc 
 contrast           estimate      SE     df t.ratio p.value
 adult - egg        -0.00790 0.01708 2233.7  -0.463  0.9906
 adult - juvenile   -0.02893 0.00625 1129.7  -4.628  <.0001
 adult - pupae      -0.00133 0.01362 2433.8  -0.097  1.0000
 adult - unknown    -0.00433 0.03103   38.5  -0.140  0.9999
 egg - juvenile     -0.02102 0.01746 2165.5  -1.204  0.7493
 egg - pupae         0.00658 0.02112 2369.4   0.311  0.9980
 egg - unknown       0.00358 0.03509   61.1   0.102  1.0000
 juvenile - pupae    0.02760 0.01340 2436.8   2.060  0.2378
 juvenile - unknown  0.02460 0.03117   39.1   0.789  0.9322
 pupae - unknown    -0.00300 0.03353   52.5  -0.090  1.0000

Degrees-of-freedom method: kenward-roger 
P value adjustment: tukey method for comparing a family of 5 estimates
```

#### Sample origin

##### Model

##### Posthoc


```
#### Enterococcus 
 contrast           estimate     SE   df t.ratio p.value
 field - laboratory  -0.0514 0.0102 2415  -5.019  <.0001

Degrees-of-freedom method: kenward-roger 

 

#### Pseudomonas 
 contrast           estimate     SE  df t.ratio p.value
 field - laboratory   -0.179 0.0232 959  -7.695  <.0001

Degrees-of-freedom method: kenward-roger
```

#### Diet study

##### Model

##### Posthoc


```
#### Wolbachia 
 contrast                  estimate     SE   df t.ratio p.value
 artificial - carnivore     -0.1528 0.0576 2432  -2.653  0.0615
 artificial - fungus         0.1752 0.0365 2421   4.795  <.0001
 artificial - herbivore      0.1366 0.0275 1981   4.972  <.0001
 artificial - unidentified   0.0217 0.0292 1922   0.743  0.9462
 carnivore - fungus          0.3280 0.0682 2436   4.813  <.0001
 carnivore - herbivore       0.2894 0.0507 2414   5.712  <.0001
 carnivore - unidentified    0.1745 0.0638 2182   2.735  0.0494
 fungus  - herbivore        -0.0386 0.0456 2383  -0.846  0.9162
 fungus  - unidentified     -0.1535 0.0467 2366  -3.284  0.0092
 herbivore - unidentified   -0.1149 0.0389 1171  -2.952  0.0267

Degrees-of-freedom method: kenward-roger 
P value adjustment: tukey method for comparing a family of 5 estimates 

 

#### Pseudomonas 
 contrast                  estimate     SE   df t.ratio p.value
 artificial - carnivore     0.17445 0.0471 2321   3.701  0.0020
 artificial - fungus        0.00549 0.0302 2436   0.182  0.9998
 artificial - herbivore     0.16685 0.0217 1048   7.696  <.0001
 artificial - unidentified -0.00168 0.0230  965  -0.073  1.0000
 carnivore - fungus        -0.16895 0.0559 2351  -3.024  0.0213
 carnivore - herbivore     -0.00760 0.0420 2428  -0.181  0.9998
 carnivore - unidentified  -0.17613 0.0510 1491  -3.455  0.0051
 fungus  - herbivore        0.16136 0.0370 2009   4.357  0.0001
 fungus  - unidentified    -0.00717 0.0378 1940  -0.190  0.9997
 herbivore - unidentified  -0.16853 0.0292  345  -5.767  <.0001

Degrees-of-freedom method: kenward-roger 
P value adjustment: tukey method for comparing a family of 5 estimates 

 

#### Leucobacter 
 contrast                   estimate     SE   df t.ratio p.value
 artificial - carnivore     0.009342 0.0214 2426   0.438  0.9924
 artificial - fungus        0.000342 0.0135 2408   0.025  1.0000
 artificial - herbivore     0.007464 0.0104 2428   0.720  0.9520
 artificial - unidentified  0.415351 0.0110 2424  37.606  <.0001
 carnivore - fungus        -0.009000 0.0252 2423  -0.357  0.9965
 carnivore - herbivore     -0.001878 0.0187 2406  -0.101  1.0000
 carnivore - unidentified   0.406010 0.0240 2436  16.946  <.0001
 fungus  - herbivore        0.007122 0.0170 2434   0.419  0.9936
 fungus  - unidentified     0.415009 0.0174 2435  23.830  <.0001
 herbivore - unidentified   0.407888 0.0150 2323  27.140  <.0001

Degrees-of-freedom method: kenward-roger 
P value adjustment: tukey method for comparing a family of 5 estimates 

 

#### Enterococcus 
 contrast                   estimate      SE   df t.ratio p.value
 artificial - carnivore     0.066828 0.01987 2431   3.363  0.0070
 artificial - fungus        0.000291 0.01255 2410   0.023  1.0000
 artificial - herbivore     0.015733 0.00963 2402   1.634  0.4759
 artificial - unidentified -0.001331 0.01025 2394  -0.130  0.9999
 carnivore - fungus        -0.066537 0.02350 2428  -2.832  0.0376
 carnivore - herbivore     -0.051096 0.01739 2407  -2.938  0.0276
 carnivore - unidentified  -0.068160 0.02226 2425  -3.062  0.0189
 fungus  - herbivore        0.015441 0.01581 2437   0.977  0.8658
 fungus  - unidentified    -0.001623 0.01620 2437  -0.100  1.0000
 herbivore - unidentified  -0.017064 0.01392 2194  -1.226  0.7361

Degrees-of-freedom method: kenward-roger 
P value adjustment: tukey method for comparing a family of 5 estimates 

 

#### Escherichia-Shigella 
 contrast                  estimate      SE   df t.ratio p.value
 artificial - carnivore    -0.08149 0.02083 2432  -3.912  0.0009
 artificial - fungus       -0.00023 0.01321 2421  -0.017  1.0000
 artificial - herbivore    -0.08507 0.00994 1982  -8.561  <.0001
 artificial - unidentified -0.01166 0.01057 1924  -1.103  0.8049
 carnivore - fungus         0.08126 0.02465 2436   3.297  0.0088
 carnivore - herbivore     -0.00359 0.01832 2414  -0.196  0.9997
 carnivore - unidentified   0.06983 0.02308 2183   3.026  0.0212
 fungus  - herbivore       -0.08484 0.01651 2383  -5.140  <.0001
 fungus  - unidentified    -0.01143 0.01690 2367  -0.676  0.9616
 herbivore - unidentified   0.07342 0.01408 1174   5.215  <.0001

Degrees-of-freedom method: kenward-roger 
P value adjustment: tukey method for comparing a family of 5 estimates 

 

#### Buchnera 
 contrast                   estimate      SE   df t.ratio p.value
 artificial - carnivore    -1.71e-01 0.00826 2409 -20.659  <.0001
 artificial - fungus        1.37e-05 0.00520 2404   0.003  1.0000
 artificial - herbivore    -2.63e-04 0.00403 2422  -0.065  1.0000
 artificial - unidentified  2.13e-04 0.00430 2423   0.050  1.0000
 carnivore - fungus         1.71e-01 0.00976 2408  17.483  <.0001
 carnivore - herbivore      1.70e-01 0.00720 2404  23.637  <.0001
 carnivore - unidentified   1.71e-01 0.00930 2418  18.363  <.0001
 fungus  - herbivore       -2.77e-04 0.00658 2412  -0.042  1.0000
 fungus  - unidentified     2.00e-04 0.00674 2413   0.030  1.0000
 herbivore - unidentified   4.76e-04 0.00588 2432   0.081  1.0000

Degrees-of-freedom method: kenward-roger 
P value adjustment: tukey method for comparing a family of 5 estimates
```

#### Sample treatment

##### Model

##### Posthoc


```
#### Salinibacter 
 contrast            estimate      SE     df t.ratio p.value
 control - treatment -0.02892 0.00868 2428.8  -3.331  0.0025
 control - unknown    0.00882 0.06192   30.6   0.142  0.9889
 treatment - unknown  0.03774 0.06243   31.6   0.604  0.8187

Degrees-of-freedom method: kenward-roger 
P value adjustment: tukey method for comparing a family of 3 estimates 

 

#### Serratia 
 contrast            estimate      SE    df t.ratio p.value
 control - treatment  0.00606 0.00542 276.0   1.118  0.5040
 control - unknown   -0.42839 0.01218  99.4 -35.165  <.0001
 treatment - unknown -0.43445 0.01313 111.4 -33.083  <.0001

Degrees-of-freedom method: kenward-roger 
P value adjustment: tukey method for comparing a family of 3 estimates 

 

#### Leucobacter 
 contrast            estimate      SE     df t.ratio p.value
 control - treatment  0.01773 0.00653 2434.1   2.716  0.0183
 control - unknown    0.00997 0.04923   30.8   0.202  0.9777
 treatment - unknown -0.00777 0.04960   31.6  -0.157  0.9866

Degrees-of-freedom method: kenward-roger 
P value adjustment: tukey method for comparing a family of 3 estimates
```

### Differentially abundant taxa - Guts


#### Insect order

##### Model

##### Posthoc


```
#### Alistipes 
 contrast                   estimate     SE   df t.ratio p.value
 blattodea - coleoptera     6.94e-02 0.0134 28.3   5.191  0.0003
 blattodea - diptera        6.94e-02 0.0129 25.1   5.387  0.0002
 blattodea - hemiptera      6.87e-02 0.0141 29.2   4.888  0.0006
 blattodea - hymenoptera    6.94e-02 0.0119 24.4   5.819  0.0001
 blattodea - lepidoptera    6.89e-02 0.0132 23.8   5.215  0.0004
 blattodea - odonata        6.92e-02 0.0221 24.3   3.132  0.0588
 coleoptera - diptera      -5.27e-06 0.0123 31.9   0.000  1.0000
 coleoptera - hemiptera    -7.27e-04 0.0135 36.6  -0.054  1.0000
 coleoptera - hymenoptera  -3.30e-06 0.0113 32.1   0.000  1.0000
 coleoptera - lepidoptera  -5.45e-04 0.0127 29.5  -0.043  1.0000
 coleoptera - odonata      -2.21e-04 0.0218 26.1  -0.010  1.0000
 diptera - hemiptera       -7.22e-04 0.0131 32.7  -0.055  1.0000
 diptera - hymenoptera      1.98e-06 0.0107 27.2   0.000  1.0000
 diptera - lepidoptera     -5.40e-04 0.0121 25.8  -0.044  1.0000
 diptera - odonata         -2.15e-04 0.0215 25.0  -0.010  1.0000
 hemiptera - hymenoptera    7.24e-04 0.0121 33.0   0.060  1.0000
 hemiptera - lepidoptera    1.82e-04 0.0134 30.4   0.014  1.0000
 hemiptera - odonata        5.07e-04 0.0222 26.5   0.023  1.0000
 hymenoptera - lepidoptera -5.42e-04 0.0111 25.0  -0.049  1.0000
 hymenoptera - odonata     -2.17e-04 0.0209 24.7  -0.010  1.0000
 lepidoptera - odonata      3.25e-04 0.0217 24.5   0.015  1.0000

Degrees-of-freedom method: kenward-roger 
P value adjustment: tukey method for comparing a family of 7 estimates 

 

#### Lactobacillus 
 contrast                   estimate     SE   df t.ratio p.value
 blattodea - coleoptera     3.67e-02 0.0262 28.1   1.399  0.7980
 blattodea - diptera        3.83e-02 0.0251 24.5   1.527  0.7268
 blattodea - hemiptera      3.84e-02 0.0276 29.2   1.389  0.8029
 blattodea - hymenoptera   -2.85e-02 0.0232 23.4  -1.230  0.8755
 blattodea - lepidoptera    3.86e-02 0.0256 22.8   1.508  0.7373
 blattodea - odonata        2.32e-02 0.0429 23.6   0.541  0.9978
 coleoptera - diptera       1.59e-03 0.0244 32.6   0.065  1.0000
 coleoptera - hemiptera     1.68e-03 0.0270 38.1   0.062  1.0000
 coleoptera - hymenoptera  -6.52e-02 0.0224 32.6  -2.911  0.0835
 coleoptera - lepidoptera   1.92e-03 0.0249 29.6   0.077  1.0000
 coleoptera - odonata      -1.35e-02 0.0425 25.7  -0.318  0.9999
 diptera - hemiptera        9.19e-05 0.0259 33.6   0.004  1.0000
 diptera - hymenoptera     -6.68e-02 0.0210 27.0  -3.177  0.0504
 diptera - lepidoptera      3.28e-04 0.0237 25.4   0.014  1.0000
 diptera - odonata         -1.51e-02 0.0418 24.4  -0.361  0.9998
 hemiptera - hymenoptera   -6.69e-02 0.0240 33.7  -2.786  0.1082
 hemiptera - lepidoptera    2.36e-04 0.0264 30.7   0.009  1.0000
 hemiptera - odonata       -1.52e-02 0.0434 26.2  -0.350  0.9998
 hymenoptera - lepidoptera  6.71e-02 0.0216 24.2   3.101  0.0630
 hymenoptera - odonata      5.17e-02 0.0407 24.0   1.271  0.8580
 lepidoptera - odonata     -1.54e-02 0.0421 23.8  -0.366  0.9998

Degrees-of-freedom method: kenward-roger 
P value adjustment: tukey method for comparing a family of 7 estimates 

 

#### Bacteroides 
 contrast                   estimate      SE   df t.ratio p.value
 blattodea - coleoptera     4.90e-02 0.01034 28.1   4.736  0.0010
 blattodea - diptera        4.90e-02 0.00988 24.5   4.958  0.0008
 blattodea - hemiptera      4.08e-02 0.01088 29.2   3.750  0.0122
 blattodea - hymenoptera    4.74e-02 0.00913 23.4   5.196  0.0005
 blattodea - lepidoptera    4.87e-02 0.01009 22.9   4.827  0.0012
 blattodea - odonata        4.90e-02 0.01690 23.6   2.897  0.0969
 coleoptera - diptera       3.31e-05 0.00959 32.6   0.003  1.0000
 coleoptera - hemiptera    -8.15e-03 0.01062 38.0  -0.768  0.9867
 coleoptera - hymenoptera  -1.53e-03 0.00882 32.6  -0.174  1.0000
 coleoptera - lepidoptera  -2.45e-04 0.00981 29.6  -0.025  1.0000
 coleoptera - odonata       2.13e-05 0.01674 25.7   0.001  1.0000
 diptera - hemiptera       -8.19e-03 0.01018 33.5  -0.804  0.9829
 diptera - hymenoptera     -1.56e-03 0.00828 27.0  -0.189  1.0000
 diptera - lepidoptera     -2.78e-04 0.00933 25.4  -0.030  1.0000
 diptera - odonata         -1.17e-05 0.01646 24.4  -0.001  1.0000
 hemiptera - hymenoptera    6.62e-03 0.00945 33.7   0.701  0.9916
 hemiptera - lepidoptera    7.91e-03 0.01038 30.7   0.762  0.9869
 hemiptera - odonata        8.17e-03 0.01708 26.2   0.479  0.9989
 hymenoptera - lepidoptera  1.29e-03 0.00853 24.2   0.151  1.0000
 hymenoptera - odonata      1.55e-03 0.01602 24.1   0.097  1.0000
 lepidoptera - odonata      2.66e-04 0.01659 23.8   0.016  1.0000

Degrees-of-freedom method: kenward-roger 
P value adjustment: tukey method for comparing a family of 7 estimates 

 

#### Desulfovibrio 
 contrast                   estimate      SE   df t.ratio p.value
 blattodea - coleoptera     4.97e-02 0.00682 27.3   7.283  <.0001
 blattodea - diptera        4.97e-02 0.00637 22.6   7.802  <.0001
 blattodea - hemiptera      4.96e-02 0.00723 29.1   6.860  <.0001
 blattodea - hymenoptera    4.97e-02 0.00582 20.3   8.530  <.0001
 blattodea - lepidoptera    4.97e-02 0.00641 20.0   7.752  <.0001
 blattodea - odonata        4.97e-02 0.01082 21.7   4.591  0.0024
 coleoptera - diptera      -1.28e-05 0.00649 35.3  -0.002  1.0000
 coleoptera - hemiptera    -1.30e-04 0.00734 42.6  -0.018  1.0000
 coleoptera - hymenoptera  -2.55e-05 0.00596 34.0  -0.004  1.0000
 coleoptera - lepidoptera  -3.30e-07 0.00654 30.2   0.000  1.0000
 coleoptera - odonata      -3.19e-05 0.01089 25.0  -0.003  1.0000
 diptera - hemiptera       -1.17e-04 0.00692 36.9  -0.017  1.0000
 diptera - hymenoptera     -1.27e-05 0.00543 26.7  -0.002  1.0000
 diptera - lepidoptera      1.24e-05 0.00606 24.7   0.002  1.0000
 diptera - odonata         -1.92e-05 0.01061 23.3  -0.002  1.0000
 hemiptera - hymenoptera    1.04e-04 0.00642 35.9   0.016  1.0000
 hemiptera - lepidoptera    1.29e-04 0.00695 32.0   0.019  1.0000
 hemiptera - odonata        9.79e-05 0.01115 25.7   0.009  1.0000
 hymenoptera - lepidoptera  2.52e-05 0.00548 22.1   0.005  1.0000
 hymenoptera - odonata     -6.46e-06 0.01030 22.5  -0.001  1.0000
 lepidoptera - odonata     -3.16e-05 0.01064 22.2  -0.003  1.0000

Degrees-of-freedom method: kenward-roger 
P value adjustment: tukey method for comparing a family of 7 estimates 

 

#### Christensenellaceae_R-7_group 
 contrast                   estimate      SE   df t.ratio p.value
 blattodea - coleoptera     3.21e-02 0.00403 27.6   7.970  <.0001
 blattodea - diptera        3.22e-02 0.00378 23.1   8.517  <.0001
 blattodea - hemiptera      3.16e-02 0.00426 29.1   7.418  <.0001
 blattodea - hymenoptera    3.22e-02 0.00347 21.2   9.283  <.0001
 blattodea - lepidoptera    3.18e-02 0.00383 20.8   8.317  <.0001
 blattodea - odonata        2.95e-02 0.00644 22.1   4.588  0.0023
 coleoptera - diptera       1.28e-04 0.00380 34.5   0.034  1.0000
 coleoptera - hemiptera    -5.15e-04 0.00428 41.2  -0.121  1.0000
 coleoptera - hymenoptera   1.33e-04 0.00349 33.6   0.038  1.0000
 coleoptera - lepidoptera  -2.75e-04 0.00385 30.0  -0.071  1.0000
 coleoptera - odonata      -2.55e-03 0.00645 25.1  -0.396  0.9996
 diptera - hemiptera       -6.44e-04 0.00405 35.8  -0.159  1.0000
 diptera - hymenoptera      4.51e-06 0.00321 26.7   0.001  1.0000
 diptera - lepidoptera     -4.03e-04 0.00359 24.8  -0.112  1.0000
 diptera - odonata         -2.68e-03 0.00630 23.5  -0.426  0.9994
 hemiptera - hymenoptera    6.48e-04 0.00376 35.2   0.173  1.0000
 hemiptera - lepidoptera    2.41e-04 0.00409 31.6   0.059  1.0000
 hemiptera - odonata       -2.04e-03 0.00660 25.8  -0.309  0.9999
 hymenoptera - lepidoptera -4.08e-04 0.00326 22.7  -0.125  1.0000
 hymenoptera - odonata     -2.69e-03 0.00612 22.9  -0.439  0.9993
 lepidoptera - odonata     -2.28e-03 0.00633 22.6  -0.360  0.9998

Degrees-of-freedom method: kenward-roger 
P value adjustment: tukey method for comparing a family of 7 estimates 

 

#### Providencia 
 contrast                   estimate     SE   df t.ratio p.value
 blattodea - coleoptera    -1.02e-02 0.0448 28.7  -0.227  1.0000
 blattodea - diptera       -2.07e-01 0.0440 26.9  -4.700  0.0012
 blattodea - hemiptera      1.82e-05 0.0469 29.3   0.000  1.0000
 blattodea - hymenoptera   -4.69e-04 0.0409 26.7  -0.011  1.0000
 blattodea - lepidoptera   -1.99e-05 0.0455 26.3   0.000  1.0000
 blattodea - odonata       -2.37e-02 0.0759 26.5  -0.312  0.9999
 coleoptera - diptera      -1.97e-01 0.0406 30.3  -4.842  0.0006
 coleoptera - hemiptera     1.02e-02 0.0437 33.0   0.233  1.0000
 coleoptera - hymenoptera   9.69e-03 0.0372 30.6   0.260  1.0000
 coleoptera - lepidoptera   1.01e-02 0.0422 29.2   0.240  1.0000
 coleoptera - odonata      -1.36e-02 0.0740 27.4  -0.183  1.0000
 diptera - hemiptera        2.07e-01 0.0429 30.8   4.827  0.0006
 diptera - hymenoptera      2.07e-01 0.0363 27.9   5.691  0.0001
 diptera - lepidoptera      2.07e-01 0.0414 27.1   5.001  0.0005
 diptera - odonata          1.83e-01 0.0735 26.8   2.492  0.2021
 hemiptera - hymenoptera   -4.87e-04 0.0397 31.2  -0.012  1.0000
 hemiptera - lepidoptera   -3.81e-05 0.0444 29.8  -0.001  1.0000
 hemiptera - odonata       -2.37e-02 0.0753 27.6  -0.315  0.9999
 hymenoptera - lepidoptera  4.49e-04 0.0381 26.9   0.012  1.0000
 hymenoptera - odonata     -2.32e-02 0.0717 26.7  -0.324  0.9999
 lepidoptera - odonata     -2.37e-02 0.0744 26.5  -0.318  0.9999

Degrees-of-freedom method: kenward-roger 
P value adjustment: tukey method for comparing a family of 7 estimates 

 

#### Wolbachia 
 contrast                   estimate    SE   df t.ratio p.value
 blattodea - coleoptera     1.00e-07 0.133 28.9   0.000  1.0000
 blattodea - diptera       -1.01e-03 0.132 28.3  -0.008  1.0000
 blattodea - hemiptera     -3.90e-01 0.139 29.1  -2.809  0.1076
 blattodea - hymenoptera   -5.46e-02 0.123 28.2  -0.443  0.9993
 blattodea - lepidoptera   -7.13e-02 0.137 28.1  -0.519  0.9983
 blattodea - odonata       -4.07e-02 0.229 28.1  -0.178  1.0000
 coleoptera - diptera      -1.02e-03 0.120 29.4  -0.008  1.0000
 coleoptera - hemiptera    -3.90e-01 0.127 30.4  -3.078  0.0593
 coleoptera - hymenoptera  -5.46e-02 0.109 29.6  -0.499  0.9987
 coleoptera - lepidoptera  -7.13e-02 0.125 29.1  -0.570  0.9972
 coleoptera - odonata      -4.07e-02 0.222 28.4  -0.183  1.0000
 diptera - hemiptera       -3.89e-01 0.126 29.6  -3.090  0.0584
 diptera - hymenoptera     -5.36e-02 0.108 28.6  -0.494  0.9987
 diptera - lepidoptera     -7.03e-02 0.124 28.3  -0.566  0.9973
 diptera - odonata         -3.97e-02 0.221 28.2  -0.179  1.0000
 hemiptera - hymenoptera    3.35e-01 0.116 29.8   2.888  0.0907
 hemiptera - lepidoptera    3.18e-01 0.131 29.3   2.430  0.2222
 hemiptera - odonata        3.49e-01 0.225 28.5   1.549  0.7139
 hymenoptera - lepidoptera -1.68e-02 0.114 28.3  -0.146  1.0000
 hymenoptera - odonata      1.39e-02 0.216 28.2   0.064  1.0000
 lepidoptera - odonata      3.06e-02 0.224 28.1   0.137  1.0000

Degrees-of-freedom method: kenward-roger 
P value adjustment: tukey method for comparing a family of 7 estimates 

 

#### Gilliamella 
 contrast                   estimate     SE   df t.ratio p.value
 blattodea - coleoptera    -2.25e-06 0.0503 28.8   0.000  1.0000
 blattodea - diptera        0.00e+00 0.0495 27.2   0.000  1.0000
 blattodea - hemiptera      0.00e+00 0.0525 29.2   0.000  1.0000
 blattodea - hymenoptera   -1.28e-01 0.0460 27.0  -2.773  0.1185
 blattodea - lepidoptera    0.00e+00 0.0512 26.7   0.000  1.0000
 blattodea - odonata       -1.07e-05 0.0855 26.8   0.000  1.0000
 coleoptera - diptera       2.25e-06 0.0455 30.1   0.000  1.0000
 coleoptera - hemiptera     2.25e-06 0.0488 32.5   0.000  1.0000
 coleoptera - hymenoptera  -1.28e-01 0.0417 30.4  -3.063  0.0614
 coleoptera - lepidoptera   2.25e-06 0.0473 29.1   0.000  1.0000
 coleoptera - odonata      -8.44e-06 0.0832 27.6   0.000  1.0000
 diptera - hemiptera        0.00e+00 0.0480 30.6   0.000  1.0000
 diptera - hymenoptera     -1.28e-01 0.0408 28.0  -3.132  0.0546
 diptera - lepidoptera      0.00e+00 0.0465 27.4   0.000  1.0000
 diptera - odonata         -1.07e-05 0.0827 27.0   0.000  1.0000
 hemiptera - hymenoptera   -1.28e-01 0.0444 31.0  -2.878  0.0914
 hemiptera - lepidoptera    0.00e+00 0.0497 29.7   0.000  1.0000
 hemiptera - odonata       -1.07e-05 0.0846 27.8   0.000  1.0000
 hymenoptera - lepidoptera  1.28e-01 0.0428 27.2   2.983  0.0766
 hymenoptera - odonata      1.28e-01 0.0807 27.0   1.582  0.6942
 lepidoptera - odonata     -1.07e-05 0.0838 26.8   0.000  1.0000

Degrees-of-freedom method: kenward-roger 
P value adjustment: tukey method for comparing a family of 7 estimates 

 

#### Snodgrassella 
 contrast                   estimate     SE   df t.ratio p.value
 blattodea - coleoptera    -3.40e-07 0.0667 28.8   0.000  1.0000
 blattodea - diptera       -5.31e-06 0.0660 27.8   0.000  1.0000
 blattodea - hemiptera      0.00e+00 0.0696 29.2   0.000  1.0000
 blattodea - hymenoptera   -1.89e-01 0.0614 27.7  -3.076  0.0623
 blattodea - lepidoptera    0.00e+00 0.0684 27.4   0.000  1.0000
 blattodea - odonata       -1.03e-04 0.1141 27.5  -0.001  1.0000
 coleoptera - diptera      -4.97e-06 0.0601 29.7   0.000  1.0000
 coleoptera - hemiptera     3.40e-07 0.0640 31.3   0.000  1.0000
 coleoptera - hymenoptera  -1.89e-01 0.0550 29.9  -3.436  0.0260
 coleoptera - lepidoptera   3.40e-07 0.0627 29.1   0.000  1.0000
 coleoptera - odonata      -1.03e-04 0.1108 28.0  -0.001  1.0000
 diptera - hemiptera        5.31e-06 0.0633 30.1   0.000  1.0000
 diptera - hymenoptera     -1.89e-01 0.0542 28.3  -3.486  0.0239
 diptera - lepidoptera      5.31e-06 0.0620 27.9   0.000  1.0000
 diptera - odonata         -9.78e-05 0.1104 27.7  -0.001  1.0000
 hemiptera - hymenoptera   -1.89e-01 0.0584 30.3  -3.234  0.0416
 hemiptera - lepidoptera    0.00e+00 0.0657 29.5   0.000  1.0000
 hemiptera - odonata       -1.03e-04 0.1125 28.2  -0.001  1.0000
 hymenoptera - lepidoptera  1.89e-01 0.0571 27.8   3.311  0.0366
 hymenoptera - odonata      1.89e-01 0.1077 27.6   1.754  0.5880
 lepidoptera - odonata     -1.03e-04 0.1118 27.5  -0.001  1.0000

Degrees-of-freedom method: kenward-roger 
P value adjustment: tukey method for comparing a family of 7 estimates
```

#### Sex

##### Model

##### Posthoc


```
#### Lactobacillus 
 contrast         estimate      SE   df t.ratio p.value
 female - male     0.02372 0.00782 1868   3.034  0.0069
 female - unknown -0.00933 0.00869  585  -1.074  0.5306
 male - unknown   -0.03305 0.00983 1201  -3.363  0.0023

Degrees-of-freedom method: kenward-roger 
P value adjustment: tukey method for comparing a family of 3 estimates 

 

#### Enterobacter 
 contrast         estimate      SE   df t.ratio p.value
 female - male    -0.01312 0.00501 1850  -2.619  0.0241
 female - unknown -0.00343 0.00603 1776  -0.569  0.8367
 male - unknown    0.00969 0.00656 1859   1.477  0.3025

Degrees-of-freedom method: kenward-roger 
P value adjustment: tukey method for comparing a family of 3 estimates 

 

#### Blattabacterium 
 contrast         estimate      SE   df t.ratio p.value
 female - male     0.01925 0.00625 1851   3.078  0.0060
 female - unknown  0.00651 0.00662  339   0.983  0.5880
 male - unknown   -0.01274 0.00768  882  -1.659  0.2219

Degrees-of-freedom method: kenward-roger 
P value adjustment: tukey method for comparing a family of 3 estimates 

 

#### Bacteroides 
 contrast         estimate      SE   df t.ratio p.value
 female - male     -0.0022 0.00304 1868  -0.724  0.7491
 female - unknown   0.0235 0.00348  880   6.766  <.0001
 male - unknown     0.0257 0.00388 1460   6.633  <.0001

Degrees-of-freedom method: kenward-roger 
P value adjustment: tukey method for comparing a family of 3 estimates 

 

#### Desulfovibrio 
 contrast         estimate      SE   df t.ratio p.value
 female - male    -0.00215 0.00262 1868  -0.820  0.6906
 female - unknown  0.02416 0.00301  927   8.040  <.0001
 male - unknown    0.02631 0.00335 1493   7.861  <.0001

Degrees-of-freedom method: kenward-roger 
P value adjustment: tukey method for comparing a family of 3 estimates 

 

#### Alistipes 
 contrast         estimate      SE   df t.ratio p.value
 female - male    -0.00501 0.00356 1866  -1.405  0.3385
 female - unknown  0.01070 0.00413 1088   2.588  0.0264
 male - unknown    0.01570 0.00458 1593   3.430  0.0018

Degrees-of-freedom method: kenward-roger 
P value adjustment: tukey method for comparing a family of 3 estimates 

 

#### Christensenellaceae_R-7_group 
 contrast          estimate      SE   df t.ratio p.value
 female - male    -0.004317 0.00148 1867  -2.925  0.0098
 female - unknown -0.000861 0.00170  969  -0.507  0.8680
 male - unknown    0.003456 0.00189 1521   1.830  0.1601

Degrees-of-freedom method: kenward-roger 
P value adjustment: tukey method for comparing a family of 3 estimates
```

#### Life stage

##### Model

##### Posthoc


```
#### Providencia 
 contrast         estimate     SE   df t.ratio p.value
 adult - egg        0.2551 0.0555 1863   4.594  <.0001
 adult - juvenile   0.0237 0.0164  908   1.448  0.4696
 adult - pupae     -0.0395 0.0214 1838  -1.846  0.2521
 egg - juvenile    -0.2313 0.0555 1861  -4.166  0.0002
 egg - pupae       -0.2946 0.0570 1851  -5.168  <.0001
 juvenile - pupae  -0.0632 0.0209 1866  -3.026  0.0134

Degrees-of-freedom method: kenward-roger 
P value adjustment: tukey method for comparing a family of 4 estimates 

 

#### Enterobacter 
 contrast         estimate     SE   df t.ratio p.value
 adult - egg        0.0306 0.0349 1859   0.877  0.8166
 adult - juvenile   0.0610 0.0105 1176   5.821  <.0001
 adult - pupae      0.0344 0.0135 1858   2.553  0.0525
 egg - juvenile     0.0304 0.0349 1856   0.872  0.8196
 egg - pupae        0.0038 0.0358 1847   0.106  0.9996
 juvenile - pupae  -0.0266 0.0131 1868  -2.023  0.1797

Degrees-of-freedom method: kenward-roger 
P value adjustment: tukey method for comparing a family of 4 estimates 

 

#### Acinetobacter 
 contrast         estimate     SE   df t.ratio p.value
 adult - egg        0.0446 0.0447 1866   0.999  0.7502
 adult - juvenile  -0.0619 0.0123  380  -5.033  <.0001
 adult - pupae     -0.0274 0.0170 1704  -1.611  0.3724
 egg - juvenile    -0.1065 0.0447 1868  -2.382  0.0809
 egg - pupae       -0.0720 0.0460 1864  -1.566  0.3983
 juvenile - pupae   0.0345 0.0167 1827   2.060  0.1668

Degrees-of-freedom method: kenward-roger 
P value adjustment: tukey method for comparing a family of 4 estimates 

 

#### Pseudomonas 
 contrast         estimate      SE   df t.ratio p.value
 adult - egg      -0.16505 0.02951 1860  -5.593  <.0001
 adult - juvenile -0.02338 0.00784  274  -2.984  0.0163
 adult - pupae    -0.02202 0.01119 1629  -1.968  0.2004
 egg - juvenile    0.14167 0.02955 1864   4.795  <.0001
 egg - pupae       0.14303 0.03042 1867   4.701  <.0001
 juvenile - pupae  0.00136 0.01103 1795   0.124  0.9993

Degrees-of-freedom method: kenward-roger 
P value adjustment: tukey method for comparing a family of 4 estimates 

 

#### Klebsiella 
 contrast         estimate      SE   df t.ratio p.value
 adult - egg       -0.0177 0.03138 1864  -0.564  0.9426
 adult - juvenile  -0.0487 0.00925  882  -5.269  <.0001
 adult - pupae      0.0224 0.01209 1835   1.856  0.2477
 egg - juvenile    -0.0310 0.03139 1861  -0.988  0.7561
 egg - pupae        0.0401 0.03222 1852   1.246  0.5978
 juvenile - pupae   0.0712 0.01181 1866   6.026  <.0001

Degrees-of-freedom method: kenward-roger 
P value adjustment: tukey method for comparing a family of 4 estimates
```

#### Sample origin

##### Model

##### Posthoc


```
#### Lactobacillus 
 contrast           estimate      SE  df t.ratio p.value
 field - laboratory  -0.0267 0.00901 369  -2.963  0.0032

Degrees-of-freedom method: kenward-roger 

 

#### Enterococcus 
 contrast           estimate     SE   df t.ratio p.value
 field - laboratory  -0.0458 0.0076 1867  -6.024  <.0001

Degrees-of-freedom method: kenward-roger 

 

#### Providencia 
 contrast           estimate     SE   df t.ratio p.value
 field - laboratory  -0.0202 0.0102 1512  -1.970  0.0490

Degrees-of-freedom method: kenward-roger 

 

#### Alistipes 
 contrast           estimate     SE  df t.ratio p.value
 field - laboratory   0.0144 0.0044 853   3.267  0.0011

Degrees-of-freedom method: kenward-roger 

 

#### Acinetobacter 
 contrast           estimate      SE   df t.ratio p.value
 field - laboratory   0.0235 0.00803 1054   2.927  0.0035

Degrees-of-freedom method: kenward-roger 

 

#### Desulfovibrio 
 contrast           estimate      SE   df t.ratio p.value
 field - laboratory   0.0278 0.00328 1051   8.479  <.0001

Degrees-of-freedom method: kenward-roger 

 

#### Bacteroides 
 contrast           estimate      SE  df t.ratio p.value
 field - laboratory   0.0128 0.00374 711   3.421  0.0007

Degrees-of-freedom method: kenward-roger 

 

#### Christensenellaceae_R-7_group 
 contrast           estimate      SE   df t.ratio p.value
 field - laboratory  0.00832 0.00183 1018   4.540  <.0001

Degrees-of-freedom method: kenward-roger 

 

#### Blattabacterium 
 contrast           estimate      SE  df t.ratio p.value
 field - laboratory  -0.0325 0.00685 246  -4.744  <.0001

Degrees-of-freedom method: kenward-roger
```

#### Diet study

##### Model

##### Posthoc


```
#### Providencia 
 contrast                  estimate     SE     df t.ratio p.value
 artificial - carnivore     0.18086 0.0376  137.5   4.814  <.0001
 artificial - herbivore     0.18320 0.0344  102.7   5.320  <.0001
 artificial - manure       -0.15202 0.1196   33.5  -1.271  0.7105
 artificial - unidentified  0.02311 0.0175 1018.4   1.318  0.6802
 carnivore - herbivore      0.00234 0.0163 1867.0   0.143  0.9999
 carnivore - manure        -0.33288 0.1221   35.0  -2.727  0.0700
 carnivore - unidentified  -0.15775 0.0389  121.9  -4.055  0.0008
 herbivore - manure        -0.33522 0.1212   34.0  -2.766  0.0646
 herbivore - unidentified  -0.16009 0.0359   93.0  -4.458  0.0002
 manure - unidentified      0.17513 0.1197   33.5   1.463  0.5927

Degrees-of-freedom method: kenward-roger 
P value adjustment: tukey method for comparing a family of 5 estimates 

 

#### Blattabacterium 
 contrast                   estimate     SE     df t.ratio p.value
 artificial - carnivore     0.036884 0.0175  121.2   2.103  0.2256
 artificial - herbivore     0.036596 0.0135   47.4   2.714  0.0667
 artificial - manure        0.038591 0.0363   36.4   1.064  0.8232
 artificial - unidentified  0.056977 0.0104  106.1   5.501  <.0001
 carnivore - herbivore     -0.000289 0.0125 1611.8  -0.023  1.0000
 carnivore - manure         0.001707 0.0384   42.6   0.044  1.0000
 carnivore - unidentified   0.020093 0.0175  110.9   1.149  0.7803
 herbivore - manure         0.001996 0.0367   36.1   0.054  1.0000
 herbivore - unidentified   0.020382 0.0135   42.9   1.514  0.5590
 manure - unidentified      0.018386 0.0362   36.1   0.508  0.9860

Degrees-of-freedom method: kenward-roger 
P value adjustment: tukey method for comparing a family of 5 estimates 

 

#### Bacteroides 
 contrast                  estimate      SE     df t.ratio p.value
 artificial - carnivore     0.01790 0.00954  111.4   1.876  0.3364
 artificial - herbivore     0.01948 0.00775   52.4   2.515  0.1029
 artificial - manure        0.02298 0.02142   34.9   1.073  0.8191
 artificial - unidentified  0.02502 0.00559  162.2   4.473  0.0001
 carnivore - herbivore      0.00158 0.00623 1723.5   0.254  0.9991
 carnivore - manure         0.00508 0.02241   39.1   0.227  0.9994
 carnivore - unidentified   0.00712 0.00957   98.8   0.744  0.9454
 herbivore - manure         0.00350 0.02172   34.7   0.161  0.9998
 herbivore - unidentified   0.00554 0.00780   46.6   0.711  0.9530
 manure - unidentified      0.00204 0.02139   34.7   0.096  1.0000

Degrees-of-freedom method: kenward-roger 
P value adjustment: tukey method for comparing a family of 5 estimates 

 

#### Desulfovibrio 
 contrast                   estimate      SE     df t.ratio p.value
 artificial - carnivore    -0.012538 0.01084  111.7  -1.156  0.7760
 artificial - herbivore    -0.012401 0.00968   74.9  -1.282  0.7032
 artificial - manure       -0.015611 0.03042   33.6  -0.513  0.9855
 artificial - unidentified -0.045687 0.00558  584.8  -8.182  <.0001
 carnivore - herbivore      0.000137 0.00540 1857.1   0.025  1.0000
 carnivore - manure        -0.003073 0.03122   35.5  -0.098  1.0000
 carnivore - unidentified  -0.033149 0.01112   97.5  -2.982  0.0291
 herbivore - manure        -0.003210 0.03085   33.9  -0.104  1.0000
 herbivore - unidentified  -0.033286 0.00999   66.8  -3.331  0.0119
 manure - unidentified     -0.030076 0.03042   33.5  -0.989  0.8585

Degrees-of-freedom method: kenward-roger 
P value adjustment: tukey method for comparing a family of 5 estimates 

 

#### Christensenellaceae_R-7_group 
 contrast                   estimate      SE     df t.ratio p.value
 artificial - carnivore    -2.87e-03 0.00592  109.8  -0.485  0.9886
 artificial - herbivore    -2.85e-03 0.00525   72.2  -0.542  0.9826
 artificial - manure       -3.49e-03 0.01631   33.7  -0.214  0.9995
 artificial - unidentified -1.76e-02 0.00309  532.0  -5.706  <.0001
 carnivore - herbivore      2.26e-05 0.00301 1853.2   0.008  1.0000
 carnivore - manure        -6.21e-04 0.01675   35.6  -0.037  1.0000
 carnivore - unidentified  -1.48e-02 0.00605   95.7  -2.437  0.1144
 herbivore - manure        -6.43e-04 0.01654   33.9  -0.039  1.0000
 herbivore - unidentified  -1.48e-02 0.00542   64.2  -2.728  0.0607
 manure - unidentified     -1.41e-02 0.01630   33.5  -0.867  0.9069

Degrees-of-freedom method: kenward-roger 
P value adjustment: tukey method for comparing a family of 5 estimates 

 

#### Wolbachia 
 contrast                  estimate     SE     df t.ratio p.value
 artificial - carnivore    -0.13564 0.0677  170.0  -2.003  0.2688
 artificial - herbivore    -0.02667 0.0628  133.2  -0.425  0.9931
 artificial - manure        0.06379 0.2356   33.5   0.271  0.9988
 artificial - unidentified -0.00944 0.0298 1300.0  -0.317  0.9978
 carnivore - herbivore      0.10897 0.0273 1864.8   3.996  0.0006
 carnivore - manure         0.19943 0.2395   34.8   0.833  0.9186
 carnivore - unidentified   0.12620 0.0704  152.5   1.792  0.3822
 herbivore - manure         0.09047 0.2383   34.1   0.380  0.9954
 herbivore - unidentified   0.01724 0.0657  121.9   0.262  0.9989
 manure - unidentified     -0.07323 0.2357   33.5  -0.311  0.9979

Degrees-of-freedom method: kenward-roger 
P value adjustment: tukey method for comparing a family of 5 estimates
```

#### Sample treatment

##### Model

##### Posthoc


```
#### Acinetobacter 
 contrast            estimate      SE     df t.ratio p.value
 control - treatment  -0.0197 0.00615 1862.7  -3.202  0.0040
 control - unknown     0.0322 0.06308   40.1   0.510  0.8669
 treatment - unknown   0.0519 0.06328   40.6   0.820  0.6930

Degrees-of-freedom method: kenward-roger 
P value adjustment: tukey method for comparing a family of 3 estimates 

 

#### Snodgrassella 
 contrast            estimate      SE     df t.ratio p.value
 control - treatment   0.0359 0.00837 1861.4   4.294  0.0001
 control - unknown     0.0544 0.13612   36.1   0.400  0.9160
 treatment - unknown   0.0184 0.13629   36.3   0.135  0.9900

Degrees-of-freedom method: kenward-roger 
P value adjustment: tukey method for comparing a family of 3 estimates 

 

#### Gilliamella 
 contrast            estimate     SE     df t.ratio p.value
 control - treatment  -0.0248 0.0079 1868.9  -3.145  0.0048
 control - unknown     0.0290 0.0931   38.4   0.311  0.9481
 treatment - unknown   0.0538 0.0933   38.7   0.577  0.8334

Degrees-of-freedom method: kenward-roger 
P value adjustment: tukey method for comparing a family of 3 estimates 

 

#### Escherichia-Shigella 
 contrast            estimate     SE     df t.ratio p.value
 control - treatment  0.00792 0.0055 1855.7   1.440  0.3204
 control - unknown   -0.24405 0.0533   41.1  -4.581  0.0001
 treatment - unknown -0.25197 0.0535   41.7  -4.713  0.0001

Degrees-of-freedom method: kenward-roger 
P value adjustment: tukey method for comparing a family of 3 estimates
```
